# Epistasis in Allosteric Proteins: Can Biophysical Models Provide a Better Framework for Prediction and Understanding?

**DOI:** 10.1101/2025.03.02.640598

**Authors:** David Ross, Drew S. Tack, Peter D. Tonner, Olga B. Vasilyeva

## Abstract

The prediction of epistasis, or the interaction between mutations, is a complex challenge impacting protein science, healthcare, and biotechnology. For allosteric proteins, the prediction of epistatic effects is further complicated by the intricate networks of conformational states and binding interactions inherent to their function. Here, we explore these issues by systematically comparing biophysical and phenomenological models to analyze mutational effects and epistasis for the lac repressor protein, LacI. Using an extensive dataset consisting of dose-response measurements for 164 LacI variants, we find that while the phenomenological Hill model provides slightly better predictive accuracy, the biophysical model fits the data more parsimoniously, with significantly less epistasis in its parameters. Our results highlight the importance of the multi-state, multi-dimensional nature of allosteric function and the potential benefits of using biophysical models for the analysis of mutational effects and epistasis.

## Introduction

How do mutations give rise to phenotypic changes? And can those changes be predicted? These fundamental questions span nearly all of biology and biotechnology, with important implications for our understanding of evolution, the progression and treatment of disease, and our ability to engineer proteins and other biological systems with useful functions. With modern DNA sequencing, synthesis, and editing technologies, we can read and write genotypes with unprecedented scale and precision. Furthermore, advances in genotype-phenotype landscape measurements (e.g., deep mutational scanning, massively parallel reporter assays) have enabled the simultaneous measurement of tens of thousands or even hundreds of thousands of genotypes along with their corresponding phenotypes [1–4]. But even these impressive new capabilities fall short of the actual need: Many important phenotypic changes only result from combinations of multiple mutations. Examples include the evolution of tryptophan synthase for optimal catalytic function in different environments [5], the engineering of fluorescent proteins with different colors [6], and the of antibiotic resistance conferred by beta-lactamase [7]. For a typical protein, comprehensive measurement even of just the two-mutation combinations is beyond current capabilities. If the combined effects of multiple mutations could be easily predicted from single-mutant data, this would not be an issue. But the effect of each mutation often depends on which other mutations are present, a phenomenon referred to as epistasis. So, the ability to predict the combined effects of multiple mutations remains an important bottleneck for modern biology and biotechnology.

Epistatic effects generally fall into two broad categories: specific epistasis, which results from direct interactions between mutations (e.g., molecular contacts); and non-specific or “global” epistasis [8–10]. With global epistasis, there is an underlying trait with additive (non-epistatic) mutational effects, and a nonlinear mapping from the additive trait to the measured phenotype. Consequently, mutational effects on the measured phenotype are non-additive (i.e., epistatic) even when the underlying trait is additive. Although it is mathematically valid to consider epistasis in the measured phenotype that arises from the nonlinear mapping, it is not very informative or parsimonious since it can obscure a simple global nonlinearity by encoding it into numerous apparent pair-wise or higher-order mutation interactions [11]. For example, in the use of mutant cycles to analyze protein folding or protein-protein interactions, it is commonly recognized that mutation effects on the dissociation constant are not additive, but that effects on the logarithm of the dissociation constant (i.e., the free energy, Δ*G*) often are additive [12, 13].

In large-scale measurements of sequence-function relationships, most of the apparent epistasis appears to be global epistasis [8–10]. So, to understand and predict mutation effects, models that accurately represent the global nonlinearities are useful since they reveal the underlying traits (i.e., parameters) for which mutation effects are most nearly additive. Furthermore, models that accurately encompass global epistasis would be expected to provide better predictive power since they would require less experimental data. For example, in a system with purely global epistasis, an accurate model of the global nonlinearities would enable prediction of most multi-mutant combinations using only data for single-mutation effects in one genetic background (e.g., the wild type). In the example of protein folding or protein-protein interactions, the most useful model of global epistasis is simple: mutation effects on the free energy of folding or binding should be additive. Deviations from this simple model (i.e., specific epistasis) can be used to identify important molecular contacts within a protein or between the protein and other molecules [12, 13].

Allosteric proteins have functions that require more than basic folding and shape-related properties such as binding. So, the choice of a model to best represent global epistasis can be more nuanced and challenging to connect to experimental data. Allosteric proteins have multiple states (i.e., conformations), usually with similar folds, and their functions involve transitions among those states. So, the effects of mutations on an allosteric protein’s function are more complex than a simple picture of mutations that stabilize or destabilize a protein fold or protein-protein interaction. If transitions among multiple states are essential to a protein’s function there should, in general, be multiple underlying traits with additive mutational effects, and global epistasis should be represented as a multi-dimensional nonlinearity (a phenomenon previously referred to as ensemble epistasis [14–16]). By analogy to the protein folding and protein-protein binding examples, it’s reasonable to assume that the underlying additive traits are related to the free energies of a protein’s different states. Based on that assumption, a model of mutational effects that is parameterized by those free energies should be more useful than a model that is not. In this article, we test this idea using data and models for mutational effects with an explicitly allosteric (and multi-state) protein: the ligand-responsive transcription factor, LacI.

The function of LacI is to regulate gene expression in response to environmental signals via binding to its cognate DNA operator. When LacI binds to its operator, it blocks (i.e., represses) transcription, so the expression level of the regulated gene(s) is low. The affinity between LacI and its operator can be modulated by binding of small molecule ligands to a distal (allosteric) site in the LacI protein. For wild-type LacI, binding of the ligand isopropyl β-D-1-thiogalactopyranoside (IPTG) reduces the LacI-operator affinity, allowing transcription. Consequently, in an environment with a high concentration of IPTG, the expression level of the regulated gene(s) is high. So, the function of LacI involves transitions between several different states: bound vs. not bound to the operator, bound vs. not bound with IPTG, etc.

There are many ways that a model of mutation effects and epistasis could be useful. But, in an effort to quantitatively compare the usefulness of different models, we start with two specific quantifiable criteria: First, a more useful model should be parameterized in a way that aligns with the global epistasis [8, 17] of the protein system, so that the model can be used to fit multi-mutant data with minimal epistasis in the model parameters. In other words, the model should represent a better choice for the “scale” on which epistasis is defined [11] and/or the “null hypothesis” [12] that mutational effects on the model’s parameters are independent and additive. Second, a more useful model should have better predictive power. Specifically, it should enable higher accuracy predictions of the effect of combined mutations (i.e., epistasis) based on measurements of single-mutation effects. We compare two different types of models against these criteria: 1) a phenomenological model, parameterized by the appropriately transformed Hill equation; and 2) a biophysical model, parameterized by the free-energy differences among the different functional states of LacI [18–23].

Previous work has shown that the biophysical model can be applied to analyze mutation effects and epistasis on the LacI dose-response: Daber et al. showed that the biophysical model could be used to explain changes in dose-response resulting from several different single mutations [18]. Chure et al. constrained the biophysical model using the results from a previous experiment [22], and analyzed the dose-response curves for both single- and double-mutant LacI variants. For double-mutants consisting of one mutation in the DNA-binding domain and a second in the ligand-binding domain (i.e., core domain), they showed that the biophysical model could be used to predict double-mutant results based on single-mutant data with the assumption of zero epistasis [20]. More recently, Morrison and Harms used the biophysical model to analyze both *in vivo* dose-response and *in vitro* binding data for a complete triple-mutant cycle in LacI (one mutation in the DNA-binding domain: M42I, and two in the ligand-binding domain: H74A and K84L) [16]. They found significant epistasis for aggregate quantities such as the effective LacI-DNA binding constant or the dose-response with IPTG, but zero epistasis for the free-energy parameters of the biophysical model – a phenomena they refer to as “ensemble epistasis” [14–16].

Here, we specifically (and quantitatively) address the question of whether the biophysical model is more *useful* than the phenomenological Hill equation model. In an attempt to be systematic, we use a much more extensive dataset than previously reported: dose-response measurements for 164 LacI variants with different combinations of mutations. Notably, we include data for the effects of a broad set of mutations, measured in multiple genetically *and phenotypically* distinct backgrounds, including the wild-type and two different backgrounds with inverted dose-response (i.e., the expression level of the regulated gene decreases with increasing ligand, IPTG). Furthermore, in the wild-type background, we also measured the effects of many multi-mutation combinations, including seven complete triple-mutant cycles. We find that the biophysical model can fit all of the data with significantly less epistasis in its parameters, but surprisingly, the phenomenological Hill model provides slightly better predictive accuracy. Beyond the quantitative criteria we use to compare the two models, we find an additional qualitative advantage for the biophysical model: it can provide more insight into the effect of mutations, particularly when mutations are applied in phenotypically distinct backgrounds (e.g., wild-type and inverted).

## Results and Discussion

To construct a dataset to assess the usefulness of different models, we used some previously reported data [24, 25] supplemented with new *in vivo* dose-response curve measurements of LacI with IPTG to give a dataset of 164 LacI variants with overlapping combinations of missense mutations (a complete list of variants and their associated data is included in Supplementary Table S1). We measured the effects of most mutations in multiple genetic backgrounds, including: the wild-type variant, a double-mutant variant with reduced response to IPTG, and two genetically distinct variants with inverted dose-response (i.e., saturated expression level of the regulated gene less than the basal expression level; Fig. 1). To choose the set of mutations and the background variants, we used a recent deep mutational scanning (DMS) dataset [24]. That DMS dataset includes the complete dose-response curves for over 60,000 LacI variants. We chose a set of 38 mutations to focus on and measured the effect of each of them in at least three different genetic backgrounds. In general, we chose those mutations based on a combination of the mutations’ locations in the LacI protein structure and their observed phenotypic effects from the DMS dataset (Table 1). The 38 mutations include multiple mutations at each of five different rheostat positions (positions where mutations can have a range of different quantitative effects on the protein’s function) [26, 27]. Most of the mutations are in different regions of the core domain of the protein, including the ligand binding pocket, the protein dimer interface, and the core-pivot region [28]. Four of the mutations are to positions in the DNA-binding domain or the linker between the DNA-binding and core domains. In addition, we measured many double- and triple-mutation combinations in different genetic backgrounds, including 53 double-mutant cycles and seven complete triple-mutant cycles.

**Figure 1.**
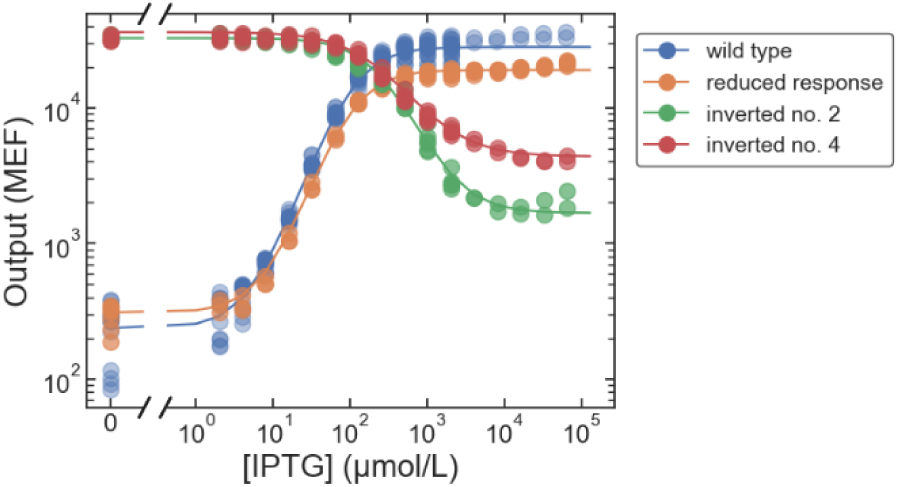
Dose response curves measured by flow cytometry for the four genetically and phenotypically distinct background LacI variants used. The reduced-response variant has two mutations: V95M and A214V. Inverted variant no. 2 has six mutations: V96E, T154I, S158R, V238D, M254I, and V264I. Inverted variant no. 4 has three mutations: A87P, V301M, and E357G. Individual points are shown for all measured replicates.

**Table 1.** Table of mutations and reasons they were chosen for this work.

| Mutations | Characteristics |
| --- | --- |
| M42I, R51C | possible epistasis from DMS dataset; M42I previously shown to stabilize the active (operator bound) state [44] |
| R51C, R51H | rheostat position; in DNA binding domain |
| R51H, Q54R | possible epistasis from DMS dataset |
| S69F, S69T | rheostat position; near ligand pocket |
| A82G, A82L, A82V | rheostat position; at dimer interface, far from ligand pocket |
| I83F, I83M | rheostat position; at dimer interface, far from ligand pocket |
| D88N, P127L | possible epistasis from DMS dataset |
| D88G, D88N | decreased $G_{\infty}$ from DMS dataset; at dimer interface, far from ligand pocket |
| F161L, F161S, F161V | rheostat position; near ligand pocket and part of core-pivot region [28] |
| A87P | single mutation that gives inverted dose-response |
| R195H, G265D, A337D | mutations making up triple-mutant band-stop |
| T68I, S70R, N125D, | near ligand pocket, expected effect on ligand affinity; |
| V80L, V136E, L296R | decreased $G_{\infty}$ and $EC_{50}$ from DMS dataset; potential decrease in $\log(k_A)$ |
| V96M, S97T | at dimer interface far from ligand pocket; expected shift in allosteric constant |
| L169P | LANTERN model [45] prediction equal to I83M, but very different position in protein structure: protein periphery far from ligand pocket or dimer interface |
| G200C | LANTERN model [45] prediction equal to F161V, but different position in protein structure: protein periphery far from ligand pocket or dimer interface |
| S279T | LANTERN model [45] prediction equal to S97T, but different position in protein structure: near, but not at the dimer interface |
| Q181H, S190C, N208Y, Q231R, V271M, Q291L | control mutations, which are nearly silent based on the DMS dataset |

For this article, we used flow cytometry measurements of the dose response curves of each variant instead of the available DMS data. Unlike the DMS dataset, which contains data for randomly applied mutations, this allowed us to examine the set of 38 mutations in targeted combinations and when applied to specific genetic backgrounds with very different phenotypes (e.g., wild-type vs. inverted dose-response). In addition, flow cytometry offers the highest possible fidelity for *in vivo* dose-response measurements, which is important for an accurate analysis of epistasis. With the random mutations of the DMS dataset, epistasis appears to be very rare, so a naïve analysis, without careful examination of possible measurement errors, can yield mostly spurious, false-positive indications of epistasis. We think this may be a general feature of large-scale DMS datasets, since their error models are often oversimplified (necessarily), to make the data analysis tractable.

To analyze the resulting data and compare the usefulness of different types of models, we fit the dose-response data using a phenomenological Hill equation model and a biophysical model previously described by Daber et al. [18], Sochor [19], and in a series of papers from the Phillips Lab [20, 22, 23, 29]. To account for uncertainty in model parameters and integrate prior information into our models, we used Bayesian inference to fit Hill equation and biophysical models to our data (Methods). To better account for multiple sources of parameter uncertainty, we included individual measurement errors as well as batch-to-batch variability in the inference. We used a model parameterization centered on the wild type, i.e., with mutational shifts of each of the parameters from the wild-type values (plus epistatic terms for multi-mutant variants). Note that although epistasis was originally used to describe non-additivity in mutation effects on an organism’s fitness, here we use the broader definition of the term to refer to non-additivity in any quantifiable phenotype. More specifically, we consider epistasis in the phenotypes that correspond to the parameters of the phenomenological and biophysical models of LacI dose-response to IPTG.

For the biophysical model, the relevant phenotypes are: the free-energy difference between the “active” and “inactive” states of LacI (Δε*_AI_*; where the active state is the state with higher affinity for the DNA operator; Δε*_AI_* is equal to the logarithm of the allosteric constant [21, 30]), the free-energy difference between LacI specifically bound to the operator and non-specifically bound to DNA (Δε*_RA_*; for LacI in the active state), and the logarithm of the binding affinities between IPTG and LacI in the active and inactive states (log(*k_A_*) and log(*k_I_*), respectively). Note that we use the same notation, parameterization, and simplifying assumptions for the biophysical model as Chure et al. [20]. Additionally, we choose the log-transformed binding affinities as the appropriate scale to correspond to free-energy differences between states of LacI [21].

For the phenomenological model, the relevant phenotypes are: the logit-transformed basal expression level and saturated expression level of the regulated gene (logit(*G*_0_/*G_max_*), and logit(*G*_∞_/*G_max_*), respectively), the logarithm of the sensitivity (log(*EC*_50_)), and the logit-transformed effective cooperativity logit(*n_eff_*/2), where *G_max_* is the maximum possible expression level from the LacI promoter-operator complex (also included as a global parameter in the biophysical mode, see Methods). We choose the log- and logit-transformed parameterization of the phenomenological model for a fair comparison, since those are expected to be the most natural transformations to a free-energy scale for phenotypes that are strictly positive (*EC*_50_) or bounded between zero and an upper limit (*G*_0_, *G*_∞_, and *n_eff_*). For *EC*_50_, this choice of parameterization is also supported by the DMS dataset, which indicates minimal epistasis for log(*EC*_50_) [24].

As a primary result, we fit all of the cytometry data simultaneously with each model, to best leverage the shared information across the set of interrelated variants. To check for consistency, we also ran similar Bayesian inference fits with sub-sets of the data, and we fit the data with two different versions of the biophysical model: a version that assumes a single copy of the lac operator per cell [20, 22], and a version that accounts for the fact that, with the plasmid system we used, there are multiple copies of the lac operator in each cell [23]. Across those different fits, the results were consistent except for variants with the mutations R51C, Q54R, and A82L (SI Fig. S1-S4). In fits with the biophysical model, the mutational effects for those mutations had complex posterior distributions (i.e., not well approximated by a multivariate normal distribution), which may have made parameter and uncertainty estimates less reliable (SI Fig. S5). So, we excluded results for variants with those three mutations from further analysis.

### Minimization of apparent epistasis

Our first criterion for a useful model is that epistasis in the model parameters is minimized. To quantitatively compare the two models with respect to that criterion, we used the Bayesian inference results for the two models to calculate the effect of mutations across different backgrounds. For the comparison, we considered a series of results with common mutations in increasingly dissimilar sets of backgrounds: first, mutations measured in double-mutant cycles; then triple-mutant cycles; and finally, mutations measured in each of the four phenotypically distinct backgrounds (wild-type, reduced response, and two inverted).

To quantify the epistasis for each model, we used a hierarchical statistical analysis to estimate the difference between the mutational effect in each background and the mean mutational effect across all backgrounds in a set. We counted a mutation as epistatic if its effect in one or more backgrounds was significantly less than or greater than the mean across backgrounds.

The apparent frequency of epistasis can depend on how experimental uncertainty is taken into account. So, to evaluate the robustness of our comparisons, we calculated the fraction of epistatic mutations using a range of different confidence thresholds to determine the significance of the difference between the individual and mean effects.

First, we analyzed the results for 53 double-mutant cycles (Fig. 2A). This case is the lowest level of background dissimilarity: each mutation characterized in two different backgrounds that differ by a single mutation. Using the lowest confidence threshold (0.5) is equivalent to ignoring experimental uncertainty entirely (e.g., using the posterior median result as truth), so the calculated epistatic fraction is 1 for both types of models. With higher confidence thresholds, the calculated epistatic fraction drops rapidly then levels off somewhat above a confidence threshold of 0.9. The epistatic fraction for the Hill equation model remains higher than for the biophysical model over the full range of confidence thresholds, indicating a higher incidence of epistasis for the Hill model. At a confidence threshold of 0.99, the epistatic fraction is about 25% for the Hill model (107 out of 424), and about 13% for the biophysical model (53 of 424), where the total number of tests, 424, is equal to the number of double-mutant cycles (53) × two mutations per cycle × the number of phenotypes (4).

**Figure 2.**
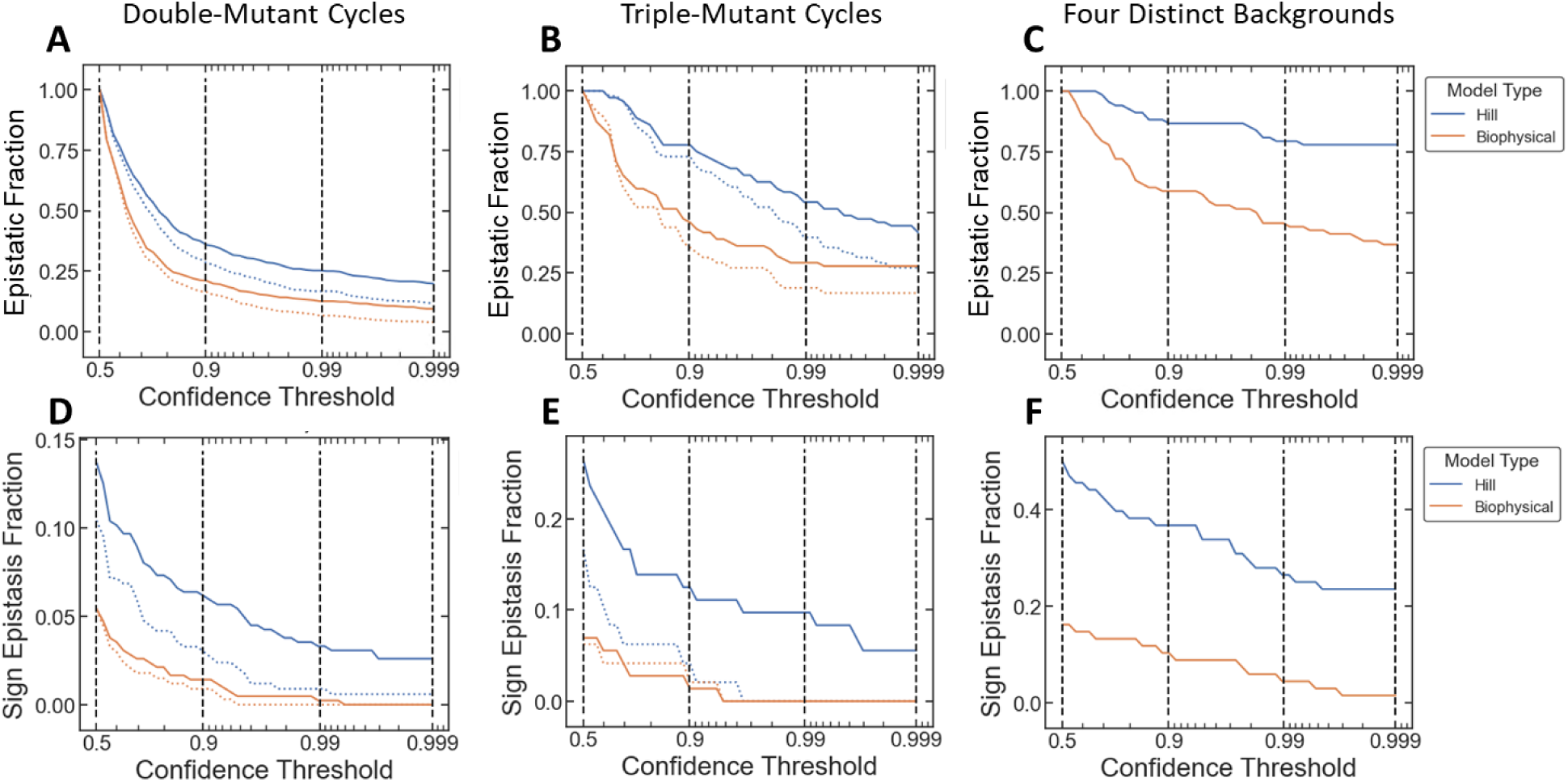
Comparison of epistasis for the Hill model (blue) and the biophysical model (orange). **A-C:** The top row shows the fraction of mutations with significant epistasis for each model using different confidence thresholds for significance. **D-F:** The bottom row shows the fraction of mutations with sign epistasis for each model using different confidence thresholds. In each plot, the solid blue and orange lines show results for all mutations in each set. In **A**, **B**, **D**, and **E**, the dotted lines show results with the V136E mutation excluded.

Next, we analyzed the results for 6 complete triple-mutant cycles (Fig. 2B). This case is an intermediate level of background dissimilarity: each mutation characterized in four different backgrounds that differ by one or two mutations. Again, as the confidence threshold increases from 0.5, the calculated epistatic fraction decreases, and the epistatic fraction for the Hill model remains higher than for the biophysical model over the full range of confidence thresholds. For both models, though, the epistatic fraction decreases more slowly and levels off at higher values than in the double-mutant-cycle analysis. At a confidence threshold of 0.99, the epistatic fraction is about twice that found for the double-mutant cycles: 54% for the Hill model (39 out of 72), and 29% for the biophysical model (21 of 72), where the total number of tests, 72, is equal to the number of triple-mutant cycles (6) × three mutations per cycle × the number of phenotypes (4).

Finally, we analyzed the results for 17 mutations, each measured in four genetically and phenotypically distinct backgrounds: the wild-type, a double-mutant variant with reduced response to IPTG, a triple-mutant with inverted dose-response, and a 6-mutant variant with inverted dose response (Fig. 1; Fig. 2C). None of the non-wild-type background variants shared any common mutations, so the mutational distance between backgrounds ranged from 2 to 9 in this case. Not surprisingly, the calculated epistatic fraction shows qualitatively similar trends as with the double- and triple-cycle results, but with epistatic fractions even higher than for the triple-mutant cycles: 79% for the Hill model (54 out of 68, at 0.99 confidence threshold), and 46% for the biophysical model (31 of 68), where the total number of tests, 68, is equal to the number of mutations (17) × the number of phenotypes (4).

As discussed below, both models had difficulty predicting the effects of the V136E mutation. So, to check the effect of that mutation on the epistatic fraction, we repeated the analysis using only the 42 double-mutant cycles and 4 triple-mutant cycles without the V136E mutation (in the analysis of the four phenotypically distinct backgrounds, none of the variants included V136E). For both the double- and triple-mutant cycles, the V136E mutation accounts for about half of the epistatic mutations (either as the mutation or in the background; dotted lines in Fig. 2A-B).

To extend our analysis to the strongest epistatic effects, we calculated the fraction of mutations with sign epistasis for the two models (Fig. 2D-F). With sign epistasis, the effect of a mutation changes sign in different backgrounds, making phenotypic prediction particularly challenging. Sign epistasis is an important determinant of landscape ruggedness [31, 32] and can indicate important molecular contacts within a protein or between the protein and other molecules [33–36]. To calculate the incidence of sign epistasis, we again used a range of confidence thresholds to determine if mutation effects were positive or negative, and we counted a mutation as having sign epistasis if its effect was significantly positive in at least one background and significantly negative in at least one other background. Across the three levels of background dissimilarity, the pattern for sign epistasis generally follows that for the epistatic fraction: The fraction of mutations with sign epistasis increases with the level of background dissimilarity, and it is consistently higher for the Hill equation model than the biophysical model. For the Hill equation model, at a confidence threshold of 0.99, the fraction of mutations with sign epistasis is 3%, 10%, and 27% for the double-mutant cycles, triple-mutant cycles, and the four phenotypically distinct backgrounds, respectively. For the biophysical model, the fraction of mutations with sign epistasis is quite low: less than 1% for the double- and triple-mutant cycles, and less than 5% for the four phenotypically distinct backgrounds (with a confidence threshold at or above 0.99).

Most examples of sign epistasis with the Hill model are for the log(*EC*_50_) phenotype (SI Fig. S6), and most involve inverted dose-response variants. The most illustrative are mutations that shift the log(*EC*_50_) one direction in the wild-type and reduced-response backgrounds, but the opposite direction in the inverted backgrounds. There are seven mutations with this type of sign epistasis for log(*EC*_50_): T68I, S69F, S70R, I83F, D88N, S97T, and F161S (SI Fig. S7). Only one of those mutations, D88N, has sign epistasis for any of the biophysical-model phenotypes (at a confidence threshold of 0.99). Four of the seven mutations are at rheostat positions: S69, I83, D88, and F161. Interestingly, the other mutations that we measured at those rheostat positions do not have sign epistasis for log(*EC*_50_) (SI Fig. S8).

These examples of sign epistasis for log(*EC*_50_) highlight a qualitative advantage of the biophysical model: it can provide insight into the mutational effects that the purely phenomenological Hill model cannot. Using just the Hill mode, we have no way to understand why mutations shift the *EC*_50_ in opposite directions in the wild-type and inverted backgrounds, or why other mutations at the same positions shift *EC*_50_ in the same direction. But, if we use the biophysical model to examine the shifts in the free-energy phenotypes resulting from those two sets of mutations, there is a pattern that provides some insight (SI Fig. S9): The set of mutations that have sign epistasis for log(*EC*_50_) generally shift Δε*_AI_* downward, to approximately zero (i.e., no free-energy difference between the active and inactive states). Also, those mutations generally shift log(*k_A_*) downward while shifting log(*k_I_*) upward, making those affinities more similar. So, from the biophysical model, it looks like these mutations are blurring the distinction between the active and inactive states. D88N is an exception to this pattern, which is not surprising since that mutation is the only one that also has sign epistasis for the biophysical model. In comparison, the set of mutations at the rheostat positions that do not have sign epistasis for log(*EC*_50_) show a different pattern of effects on the free-energy phenotypes. They also shift Δε*_AI_* downward, but not as much (on average). But, with the exception of S69T and D88G, this second set of mutations shifts both log(*k_A_*) and log(*k_I_*) upward, indicating a generally weaker affinity for the IPTG ligand. So, in contrast with the Hill model, the biophysical model handles these different sets of mutations naturally, with minimal epistasis, as having different effects on the functionally relevant states of the protein.

For the double-mutant cycle results, we also analyzed the data to determine the fraction of mutations with reciprocal sign epistasis: where the effect of both mutations changes sign when they are combined. At a confidence threshold of 0.99, there are no examples of reciprocal sign epistasis for the biophysical model, and only two double-mutant cycles (out of 212) have reciprocal sign epistasis for the Hill model (SI Fig. S10). Those two double-mutant cycles are: 1) A87P and V301M in the wild-type background; and 2) I83M and S97T in the V136E background. Both of those double-mutant cycles include variants with inverted dose-response, and similar to the sign epistasis results, both involve a change in sign of the mutational effect on the log(*EC*_50_) phenotype (SI Fig. S11).

Reciprocal sign epistasis is sometimes interpreted as an indication of molecular contact or interactions between the mutated residues or between those residues and other molecules [33, 34, 36] (e.g., for LacI, IPTG or the DNA operator). However, specific molecular interactions would be expected to result in strong epistasis for both models, not just the Hill model as with the examples here. Nevertheless, to see if there are any correlations with the protein structure or molecular contacts, we used the published crystal structures [37] to examine the two reciprocal-sign-epistatic double-mutant cycles. For each cycle, we calculated distances between the two mutations in the cycle and between each of those mutations and the DNA operator and IPTG molecule. The two residues for the first double-mutant cycle (A87, V301) have alpha carbons that are about 6.5 Å apart, so a direct molecular interaction is possible. For the second double-mutant cycle, however, the residues (I83, S97) have alpha carbons that are about 14 Å apart, ruling out any direct molecular interaction. Also, all of the residues from both double-mutant cycles have alpha carbons at least 10 Å from the IPTG molecule and at least 20 Å from the DNA operator.

To increase the statistical power of the structural analysis, we extended it to the double-mutant-cycle results for sign epistasis (reciprocal or otherwise; Fig. 2D). To assess the statistical significance, we used the two-sample Kolmogorov-Smirnov test to compare the distributions of molecular distances for double-mutant cycles with and without sign epistasis. For the distance between mutations in each cycle and the distance between those mutations and the IPTG ligand, there is no statistically significant difference. However, the residues where mutations have sign epistasis are significantly closer to the DNA operator than other residues in the double-mutant cycle data. The residues with sign epistasis are all at least 20 Å away from the closest atoms on the DNA chain though, so this difference does not indicate direct molecular contacts. Instead, all but one of the sign-epistasis residues are in the n-terminal core sub-domain (the portion of the core closer to the DNA molecule), whereas none of the non-sign-epistasis residues are in that sub-domain. So, these examples of sign epistasis are correlated with the protein structure, but not in a manner that is related to specific molecular contacts or interactions.

### Predictive accuracy from different models, assuming zero epistasis

Our second criterion for a useful model is that it can provide better predictive accuracy. To test the two models against that criterion, we used data for single mutations in the wild-type background to predict the mutational effects in other backgrounds. Specifically, we used the Bayesian inference methods to fit a sub-set of the data consisting of cytometry results for single mutations measured only in the wild-type background, plus results for four phenotypically distinct background variants: the wild-type, the low-response background, and the two inverted backgrounds. We then used the results of those fits, with the assumption of zero epistasis in the model parameters, to predict the dose-response of multi-mutant LacI variants, including double-mutants and triple-mutants in the wild-type background and the single mutations in the three non-wild-type backgrounds.

To quantitatively evaluate the predictive accuracy of each model, we compared the zero-epistasis predictions with the experimental results for the multi-mutant variants (Fig. 3). We used multiple metrics to evaluate the predictive accuracy, including the Pearson correlation coefficient between the predicted and measured values, the logarithmic root-mean-square error, and the fraction of predictions within 1.5-fold, 2.5-fold, and 5-fold of the measured values (Table 2). Contrary to our expectation that the biophysical model would provide better predictive accuracy, the Hill model performed slightly better based on almost every metric. The biophysical model only had better performance for the fraction of predictions within 1.5-fold.

**Figure 3.**
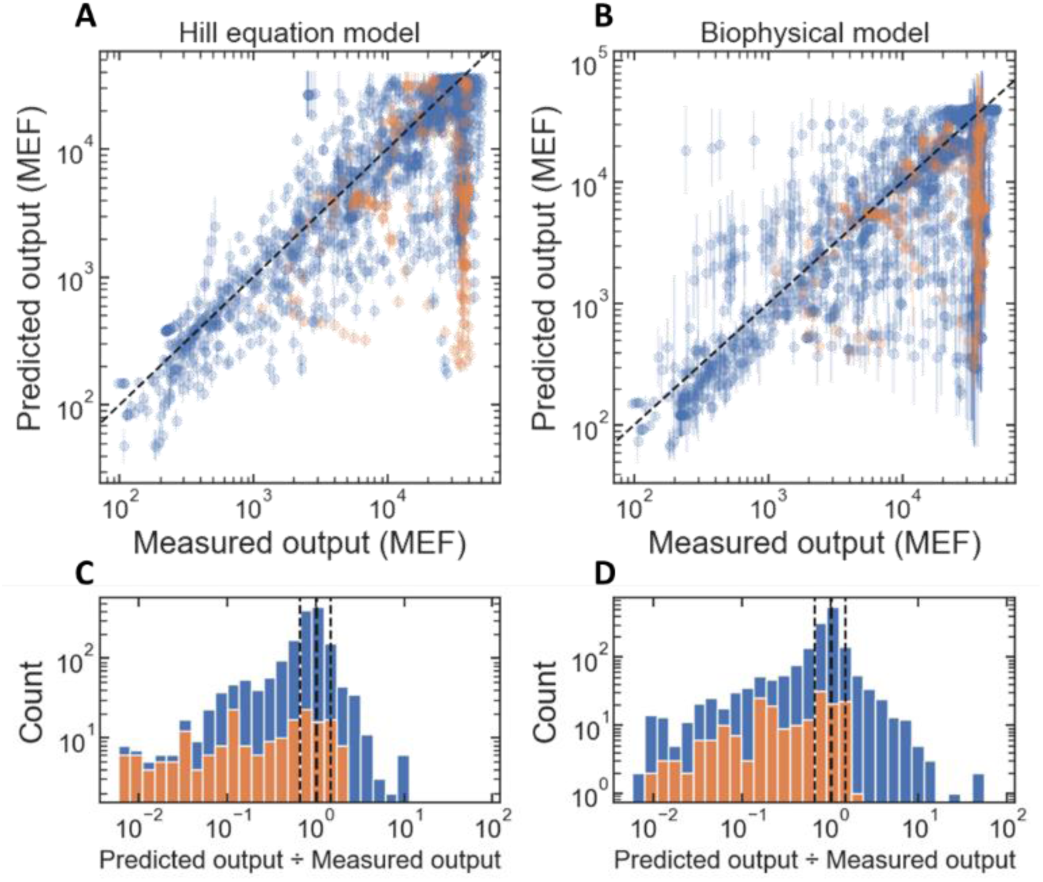
Evaluation of the predictive accuracy of the Hill model and biophysical model. **A-B:** Scatter plots of the predicted vs. measured dose-response output for the multi-mutant LacI variants. Orange points indicate results for variants with the V136E mutation. The error bars indicate ± one standard deviation of the posterior distribution for each predicted point (on the log scale). The predictions were made using data for single mutations in the wild-type background, plus data for the four phenotypically distinct backgrounds (wild-type, reduced-response, and two inverted backgrounds). The multi-mutant variants used for the evaluation consisted of double- and triple-mutant combinations in the wild-type background, plus the single mutations in the three non-wild-type backgrounds. **C-D:** Histograms of the prediction error. The blue histogram bars are results for all of the variants; the orange histogram bars are for results for variants with the V136E mutation. The vertical dashed lines indicate the range of predicted values within 1.5-fold of the measured values.

**Table 2.** Performance metrics for the predictive accuracy of the Hill equation and Biophysical models. The data used to calculate the performance metrics is plotted in Fig. 2. The log-RMSE metric is the root-mean-square error of the base-10 logarithm of the predicted dose response. The fold-RMSE, equal to 10^log-RMSE^, indicates the typical fold-accuracy of the predictions. For the Pearson correlation coefficient and the log-RMSE, the values listed are the posterior mean ± one standard deviation.

|  | model type | number of data points | Pearson R | log-RMSE | fold-RMSE | fraction within 1.5-fold | fraction within 2.5-fold | fraction within 5-fold |
| --- | --- | --- | --- | --- | --- | --- | --- | --- |
| all variants | Hill equation | 1706 | **0.785 ± 0.010** | **0.500 ± 0.017** | **3.160** | 0.563 | **0.770** | **0.860** |
|  | Biophysical | 1706 | 0.725 ± 0.029 | 0.568 ± 0.038 | 3.696 | **0.570** | 0.744 | 0.843 |
| variants without V136E | Hill equation | 1513 | **0.862 ± 0.007** | **0.399 ± 0.015** | **2.506** | **0.605** | **0.810** | **0.902** |
|  | Biophysical | 1513 | 0.766 ± 0.029 | 0.531 ± 0.040 | 3.395 | 0.601 | 0.777 | 0.874 |
| Inverted variants | Hill equation | 770 | **0.566 ± 0.017** | **0.523 ± 0.020** | **3.333** | **0.536** | **0.686** | **0.809** |
|  | Biophysical | 770 | 0.536 ± 0.043 | 0.672 ± 0.063 | 4.7 | 0.534 | 0.678 | 0.783 |

For both models, most of the prediction error is for variants with measured dose-response near the maximum value (≈ 3×10^4^ molecules of equivalent fluorophore, MEF), but a lower predicted dose-response. Looking more closely at those variants, many of them have an inverted dose-response that was not predicted by either model (e.g., SI Fig. S12A-D). Two additional double-mutant variants have a band-stop dose-response phenotype that is inconsistent with the sigmoidal form of the dose response for both models (SI Fig. S12E-F). Those inverted and band-stop variants have been reported previously [24, 25], and Tack et al. [25] noted that most of them include the same mutation: V136E. Interestingly, the predictions for those inverted and band-stop variants are generally accurate at high IPTG concentrations, but not at lower IPTG concentrations (SI Fig. S12). To test the effect of the V136E mutation on the predictive accuracy comparison, we recalculated the accuracy metrics using only results for variants without the V136E mutation (blue data points in Fig. 3A-B). The accuracy metrics improved for both model types, but the Hill equation model still outperformed the biophysical model (by every metric in this case; Table 2).

In our comparison of the two models across increasingly dissimilar sets of backgrounds, we found that the biophysical model has clear advantages for the interpretation of mutational effects and minimization of epistasis across phenotypically dissimilar backgrounds (e.g., wild-type and inverted). So, we repeated the predictive-accuracy comparison, with a focus on prediction of mutational effects in phenotypically distinct backgrounds. To that end, we tested how well each model could predict the effect of mutations in inverted backgrounds using data for single mutations in the wild-type background. For the same prediction task that we considered above, prediction of the values along the dose-response curves, the Hill model still outperforms the biophysical model (Table 2, bottom two rows). But, since the strongest epistatic effects are for the log(*EC*_50_) phenotype of the Hill model, we also compared the models with a different prediction task: predicting the *EC*_50_ resulting from mutations applied to inverted backgrounds, again using just single-mutant effects in the wild-type background. In this case, the biophysical model, with a Spearman correlation coefficient of 0.44, clearly outperformed the Hill model, with a Spearman correlation coefficient of -0.04 (SI Fig. S13). This again highlights the ability of the biophysical model to parsimoniously capture mutational effects across a broader range of backgrounds.

## Conclusions

As observed by the statistician George Box, “All models are wrong but some are useful” [38, 39]. Here, we quantitatively compare the usefulness of two different kinds of models, a biophysical model and a phenomenological model, applied to the problem of understanding and predicting mutational effects and epistasis. For our first criterion for model usefulness (minimal epistasis), the biophysical model is clearly better as it can be used to capture mutational effects with minimal epistasis. Under the second criterion for model usefulness (predictive accuracy), the Hill model is slightly better for the most general predictive tasks, but the biophysical model can be more accurate when predicting mutational effects across dissimilar backgrounds, e.g., wild-type and inverted LacI variants. Beyond our quantitative criteria for usefulness, each model has different qualitative advantages and disadvantages: The biophysical model can provide insight into mechanisms underlying mutational effects (e.g., how mutations affect the protein’s functional states). The parameters of the Hill model are directly related to the experimental observations, so it is easy and intuitive to make connections between that model and the data for quantitative interpretation of the directly observed phenotypes. In contrast, the biophysical model has potentially ambiguous connections to the data: observed mutational effects can typically be explained equally well by changes to multiple different biophysical parameters [20, 40]. This ambiguity can be mitigated by measuring the effects of mutations in different backgrounds, as demonstrated here for allosteric dose-response, and by Faure et al. for a protein folding-binding system [41]; or with additional experiments to independently measure one or more of the biophysical parameters [18, 20, 22]. Ultimately, the most useful approach is probably to combine results from both types of models. For example, with our analysis of sign epistasis, we used the Hill model to identify categories of mutations based on their apparent epistasis for the directly observed phenotypes, then we used the biophysical model to gain insight into the mechanism underlying those different mutation categories (i.e., how they affect the free energies of different protein states).

Finally, our results highlight another important consideration when using experimental data to quantify the prevalence of epistasis: The apparent frequency of epistatic interactions can strongly depend on the details of the analysis. In particular, the apparent fraction of mutations with epistasis can vary over nearly the full range from zero to one, depending on how the experimental uncertainty is accounted for in the analysis (e.g., the different confidence thresholds in Fig. 2A-C). In pointing this out, we don’t mean to say that meaningful estimates of the epistatic fraction cannot be obtained, just that researchers in the field should take care to examine the robustness of their conclusions over different treatments of experimental uncertainty. As with any problem that balances the trade-off between false positives and false negatives, the “correct” approach depends on the goals of a given study.

More fundamentally, however, our results demonstrate that the epistatic fraction can depend on the choice of model used with the data (i.e., the “scale” associated with the null hypothesis that mutational effects are independent and additive). This point has been made many times in previous reports [11], but is still worth repeating, particularly as large-scale DMS datasets become increasingly accessible.

Our use of the logit and log transformations for the parameters of the Hill model is similar to the nonlinear transformations often used to minimize the effects of global (i.e., non-specific) epistasis [6, 9, 11, 42, 43], since it similarly accounts for the upper and lower bounds of the measurements. So, the large difference in epistatic fractions between the Hill and biophysical models might be unexpected. Similar to the conclusions of Sailor and Harms [14] and Morrison et al. [15], we think this large difference is due to the multi-state, multi-dimensional nature of the allosteric function of LacI: The logit and similar transformations are one-dimensional corrections applied to a single measured phenotype. On the other hand, the non-linear transformations between the Hill and biophysical models are inherently multidimensional, since each observed phenotype for the Hill model depends on a mixture of all four free-energy phenotypes of the biophysical model. In short, our results suggest that for complex protein functions such as allosteric signaling, the most appropriate scale for parsimonious interpretation of mutational effects and epistasis will require multi-dimensional transformations to account for multi-dimensional global epistasis (i.e., ensemble epistasis). We think this inherent multi-dimensional nature of protein function will become increasingly important as more mutational studies extend beyond simple protein folding and binding.

## Methods

### Strain, plasmids, and cell culture conditions

All reported measurements were done with *E. coli* strain MG1655Δ*lac* [46]. Briefly, strain MG1655Δ*lac* is a complete lactose-operon knockout constructed from *E. coli* strain MG1655 (ATCC #47076) by replacing the lactose operon with the bleomycin resistance gene from *Streptoalloteichus hindustanus* (*Shble*).

For dose-response measurements of LacI variants, the previously described pVER plasmid was used [24]. Plasmid pVER contained two expression cassettes, one for constitutive expression of LacI, and a second in which LacI and the lactose operator (*lacO*) regulate the expression of Enhanced Yellow Fluorescent Protein (YFP). For the measurements, each LacI variant was chemically synthesized (Twist Biosciences), inserted into pVER (by restriction cloning with XhoI and AscI), and transformed into *E. coli* strain MG1655Δ*lac* for flow cytometry measurements. The annotated sequence of the pVER plasmid, with wild-type LacI, is included here as Supplementary File S1, in GenBank format.

Non-fluorescent control measurements were also made with MG1655Δ*lac* cells containing a plasmid similar to pVER but lacking the YFP gene (pAN-1201 from Nielsen et al. [47]).

The maximum regulated gene expression from the lactose operator, *G_max_*, is the expression level from the operator when it is not bound by LacI. To measure *G_max_* we used the pVER plasmid with a LacI variant that had the n-terminal 60 amino acids (the DNA binding domain) deleted.

All cultures were grown in a rich M9 media (3 g/L KH_2_PO_4_, 6.78 g/L Na_2_HPO_4_, 0.5 g/L NaCl, 1 g/L NH_4_Cl, 0.1 mmol/L CaCl_2_, 2 mmol/L MgSO_4_, 4% glycerol and 20 g/L casamino acids) supplemented with 50 μg/mL kanamycin. Cultures were grown in 96-well plates with automated microbial culture and measurement system following a previously described protocol [48]. Briefly, cultures of *E. coli* containing each plasmid were started from glycerol stock scrapings in M9 media (5 mL of culture in 14 mL snap-cap culture tubes). Cultures were grown overnight (12–20 h) at 37 °C with shaking at 300 rpm. Cultures were then loaded into the automated microbial culture system, where the following protocol was run:

1. Dilute cultures 10x from the culture tubes into 96-well plates (4titude part number 4ti-0255, square wells, 1.1 mL per well maximum volume. 0.5 mL per well culture volume).
2. Apply a gas-permeable seal to the 96-well plate (4titude part number 4ti-0598).
3. Incubate for 12–13.5 h in a plate reader (Biotek Neo2SM).

a. 37 °C, with double-orbital shaking at 807 cycles per minute.
4. Remove the gas permeable seal and dilute the culture from each well in the 96-well plate 50x into a second 96-well plate prepared with M9 media and different concentrations of IPTG.
5. Apply a gas-permeable seal to the second 96-well plate.
6. Incubate for 160 min in a plate reader, at 37 °C, with double-orbital shaking at 807 cycles per minute.

a. This allows the cultures to grow up to mid-log phase with induction.
7. Remove the gas permeable seal and dilute the culture from each well in the second 96-well plate 10x into a third 96-well plate prepared with M9 media and the same concentrations of IPTG.
8. Apply a gas-permeable seal to the third 96-well plate.
9. Incubate for 160 min in a plate reader, at 37 °C, with double-orbital shaking at 807 cycles per minute.

a. This allows the cultures to grow up to mid-log phase a second time with induction.
10. Remove the gas permeable seal and dilute the culture from each well in the third 96-well plate 40x into a round-bottom 96-well plate (Falcon part no. 351177) prepared with phosphate-buffered saline (PBS) with 170 µg/ml chloramphenicol to halt the translation of YFP.

a. 5 µL of each culture plus 195 µL PBS

### Flow cytometry measurements

All of the data used for this Article was collected with flow cytometry to measure the dose-response curves of different LacI variants. Samples prepared in PBS with chloramphenicol were measured on an Attune NxT flow cytometry with autosampler using a 488 nm excitation laser and a 530 nm ± 15 nm band-pass emission filter.

An automated gating algorithm was used to discriminate cell events from non-cell events and singlet events from multiplet events using blank samples that were measured with each batch of cell measurements [49].

An additional automatic unsupervised gating procedure, similar to that described by Razo-Mejia et al. [22], was then used to select the region of the side-scatter vs forward-scatter plots containing the highest density of singlet cell events. The additional automatic gating was set to retain 30% of the singlet cell events. Fluorescence calibration beads (Spherotech, part no. RCP-30-20A) were also measured with each flow cytometry plate to enable calibration of the flow cytometry data to molecules of equivalent fluorophore [50–52].

All cytometry data are included in Supplementary Table S1.

### Models for LacI Dose-Response

The equation used to fit the dose-response data for the Hill model is:

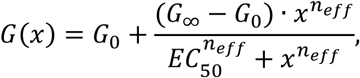

where *x* is the concentration of IPTG, *G*(*x*) is the regulated gene expression output, *G*_0_ is the basal expression level, *G*_∞_ is the saturated gene expression level, *EC*_50_ is the concentration required for an output mid-way between *G*_0_ and *G*_∞_, and *n_eff_* is the Hill coefficient.

The equation used to fit the dose-response data for the multi-operator biophysical model (the one used for most of the Results) is from Razo-Mejia et al. [22]:

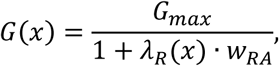

where *G_max_* is the maximum possible expression level from the LacI promoter-operator complex, and *w_RA_*

is given by:

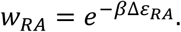

Δε*_RA_* is the free-energy difference between LacI specifically bound to the operator and non-specifically bound to DNA, β is the inverse temperature, and the LacI fugacity, λ*_R_*, is given by:

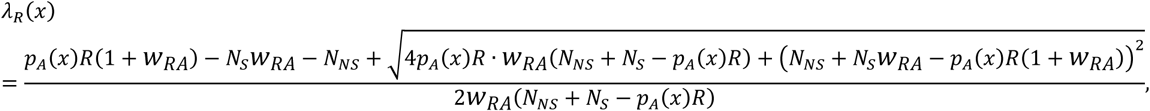

where R is the number of LacI dimers per cell, *N_S_* is the copy number of the *lacO* operator per cell, *N_NS_* is the number of non-specific DNA binding sites (≈ the length of the *E. coli* genome), and the probability that LacI is in the active state, *p_A_*(*x*), is given by:

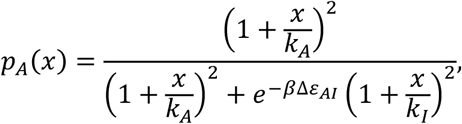

where Δε*_AI_* is the free-energy difference between the “active” and “inactive” states of LacI, and *k_A_* and *k_I_* are the binding affinities between IPTG and LacI in the active and inactive states, respectively.

The equation used to fit the dose-response data for the single-operator biophysical model (used to check consistency of results) is also from Razo-Mejia et al. [22]:

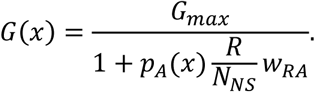

### Bayesian parameter estimation

Fits to the dose-response data were performed using Bayesian inference with Markov chain Monte Carlo (MCMC) using the cmdstanpy interface to Stan [53, 54]. Files specifying the Stan models are included in Supplementary File S2. All fits were run with four independent chains.

Fits were performed for three different types of sub-sets of the data:

First, fits were performed for each model type (Hill, single-operator biophysical and multi-operator biophysical) using all of the dose-response data simultaneously. These fits were run with 3000 iterations (1500 warmup, 1500 sampling), using a model parameterization that included terms for mutational effects and epistasis. The time to run these fits took from about 10 days for the Hill model to over two weeks for the multi-operator biophysical model. The Gelman-Rubin *R̂* diagnostic was used to check convergence of each fit. The *R̂* score was generally less than 1.05 for all model parameters except those related to the mutations R51C, Q54R, and A82L, which as noted in the Results, had strangely shaped posterior distribution that made sampling more difficult (SI Fig. S5).

Second, to check the reliability of the fits with all of the data, fits were performed using several different sub-sets of the data: for each mutation in Table 1, fits were run using data for all variants with that mutation plus the four phenotypically distinct background variants (wild-type, reduced-response, and two inverted). These fits were run with 6000 iterations (3000 warmup, 3000 sampling), using a model parameterization that included terms for mutational effects and epistasis. The time to run these fits ranged from about 10 hours to about 30 hours each, though they were typically run several at a time in parallel. As with the fits using all of the data, the *R̂* score was generally less than 1.05 for all model parameters except those related to the mutations R51C, Q54R, and A82L.

Third, to test the predictive accuracy of each model type, fits were performed using a sub-set of the data consisting of results for the four phenotypically distinct background variants plus variants with single mutations in the wild-type background. These fits were run with 3000 iterations (1500 warmup, 1500 sampling), using a model parameterization that only included terms for mutational effects (but not epistasis). The time to run these fits was between one and two days. The *R̂* score was less than 1.05 for all model parameters in this case.

With the biophysical models, ambiguities in the relationship between the free-energy parameters and the dose-response make it difficult to estimate those parameters from a single dose-response measurement [20]. Previous work mitigated this ambiguity with additional experiments to independently measure one or more of the biophysical parameters [18, 20, 22]. Here, the ambiguity was partially mitigated by fitting to many variants simultaneously as detailed above. This was particularly helpful for mutations that were measured in both the wild-type and one or both inverted backgrounds. In addition, the ambiguity was primarily mitigated in this work by using the results from previous publications as informative priors on the free-energy parameters for wild-type LacI [20, 22, 55]. More specifically, for the wild-type free-energy parameters, we used normal distribution priors with the following means and standard deviations:

| parameter | prior mean | prior standard deviation | posterior mean | posterior standard deviation |
| --- | --- | --- | --- | --- |
| $\Delta\epsilon_{AI}$ | 4.5 | 0.1 | 4.659 | 0.064 |
| $\Delta\epsilon_{RA}$ | -15.2 | 0.003 | -15.2 | 0.003 |
| $\log(k_A)$ | 4.93 | 0.092 | 4.657 | 0.040 |
| $\log(k_I)$ | -0.636 | 0.039 | -0.494 | 0.032 |

The table also lists the posterior mean and standard deviation for each parameter (from the fit using the multi-operator biophysical model with all of the cytometry data). For Δε*_AI_* and Δε*_RA_*, the posteriors are not significantly different from the priors, indicating that, for those parameters, the data collected here is consistent with the prior information from previous publications [20, 22, 55]. For log(*k_A_*) and log(*k_I_*), the posteriors ***are*** significantly different from the priors in a statistical sense, but those differences are relatively small in absolute terms: They imply a 1.3-fold difference in *k_A_* and a 1.15-fold difference in *k_I_*, which can reasonably be attributed to systematic differences in the way the experiments were done between the current and previous work (e.g., plasmid-borne vs. genomically integrated expression cassettes).

All of the Bayesian inference models included parameters for mutational effects, with zero-mean, normal distribution priors.

With the biophysical models, we used priors with a standard deviation of 10 for the mutational effects on Δε*_AI_* and, log(*k_A_*), and log(*k_A_*). For mutational effects on Δε*_RA_*, we used priors with a standard deviation of 10 for mutations to residues in the DNA binding domain or the hinge connecting the DNA binding and core domains (residues 1-61); for mutations outside those regions, we used priors with a standard deviation of 0.5. Note that this imposes a prior assumption that mutations far from contacting the DNA operator probably have only a small effect on the LacI-operator affinity. However, it allows sufficient flexibility for this assumption to be overridden by the experimental data. In this respect, it is a less stringent prior assumption than has been applied in previous work where mutational effects on Δε*_RA_* were set equal to zero for mutations in the core domain [18, 20]. In fact, in this work, of 49 mutations in the dataset at residues far from DNA contacts, 15 were found to have posterior values for the mutational effect on Δε*_RA_* that were significantly different from zero (with 0.99 confidence), though the mean posterior mutational effect is greater than 1 for only two of them.

With the Hill models, we used a zero-mean normal prior for the mutational effect on all parameters (logit(*G*_0_/*G_max_*), logit(*G*_∞_/*G_max_*), log(*EC*_50_), and logit(*n_eff_*/2)).

The Bayesian inference models with epistasis (see above) included parameters for the total epistasis for each multi-mutant LacI variant. For those parameters, we used a continuous version of a spike-and-slab prior: a mixture of two zero-mean normal distributions: one narrow and the other wide. This imposed a prior assumption with a preference for zero epistasis, but with broad flexibility if the data are inconsistent with that assumption. The mixture weight for the narrow part of the prior was 0.3. For the biophysical models, the narrow and wide components of the prior had standard deviations of 1 and 10, respectively. For the Hill model, the narrow and wide components of the prior had standard deviations of 0.5 and 5, respectively.

Within each type of model (Hill, single-operator biophysical and multi-operator biophysical), there are some parameters of the dose-response equations that we assumed were unaffected by mutations. We refer to these as “global” parameters. For the Hill model, and the single-operator biophysical model there is only one global dose-response parameter: *G_max_*. We used a log-transformed parameterization for *G_max_*, with a prior for log_10_(*G_max_*) informed by previous experience with the pVER plasmid system (mean 4.6, standard deviation 0.2).

For the single-operator biophysical model, the dose-response also depends on the ratio, *R*/*N_NS_*, but that ratio is only present in the dose-response equation when multiplied by the factor *e*^−*β*Δε*RA*^. So, *R* and *N_NS_* cannot be determined as separate parameters from fits to the dose response. Accordingly, we set them as constants in the single-operator biophysical model: *R* = 200, *N_NS_* = 4.6×10^6^ (the length of the *E. coli* genome).

For the multi-operator biophysical model, the global dose-response parameters are: *G_max_*, *R*, and *N_S_*, and we set the number of non-specific DNA binding sites as a constant, *N_NS_* = 1.38×10^7^ (three times the length of the *E. coli* genome to account for the multi-genome copy number of *E. coli* in exponential growth [56]). We used a log-transformed parameterization for *R* and *N_S_*. For log_10_(*R*), we used a broad normal prior with mean of 1.9 and standard deviation of 2. For log_10_(*N_S_*), we used a normal prior informed by previously published results for the copy number for a plasmid similar to pVER [57] (mean 0.954, standard deviation 0.1).

Each model also included parameters related to the measurement process, including the non-fluorescent background signal, and batch effects to account for small variations in the background and gain for measurements made on different days.

In addition to the flow cytometry for the LacI variants described in the Results, measurements were also made using two control plasmids: 1) a non-fluorescent control plasmid (pAN-1201) to provide direct data to estimate the measurement background and its variability, and 2) a plasmid containing a LacI variant with the DNA binding domain deleted (LacI-WT(N-Δ60)) to give direct data to estimate *G_max_* and its variability. Data for these control results were included as inputs to every fit.

## Appendix A. Supplementary material

The following are the Supplementary material to this article:

Supplementary Figures.

Supplementary Table S1. This table contains all cytometry data used for this work.

Supplementary File S1. The annotated sequence of the pVER plasmid, with wild-type LacI, in GenBank format.

Supplementary File S2. Files specifying the statistical models used with Stan for model fitting.

## Data availability

All processed cytometry data (mean fluorescence per cell) are included as Supplementary Table S1. Raw cytometry data will be made available on request.

## Disclaimer

Certain commercial equipment, instruments, or materials are identified to adequately specify the experimental procedure. Such identification implies neither recommendation or endorsement by the National Institute of Standards and Technology nor that the materials or equipment identified are necessarily the best available for the purpose. The views expressed in this publication are those of the authors and do not necessarily represent the views of the U.S. Department of Commerce or the National Institute of Standards and Technology.

## Acknowledgements

We would like to thank Liskin Swint-Kruse for insightful comments on the manuscript.

## Declaration of generative AI and AI-assisted technologies in the writing process

During the preparation of this work the authors used a local instance of LLaMA 3.3 to suggest sentences for the abstract and title. After using this tool, the authors reviewed and edited the content as needed and take full responsibility for the content of the publication.

