## Supplementary Figures for "Epistasis in Allosteric Proteins: Can Biophysical Models Provide a Better Framework for Prediction and Understanding?"

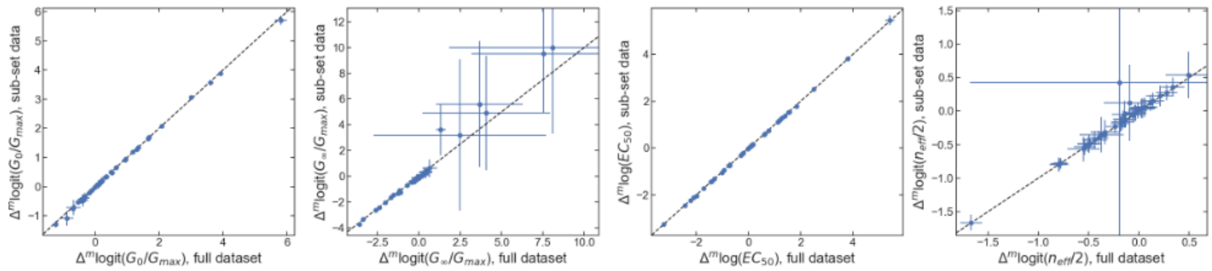


**Figure S1**. Comparison between Hill model fit results using the full dataset (x-axis) and sub-sets of the data for each mutation (y-axis). Plotted points are the posterior mean and error bars indicate ± one posterior standard deviation.

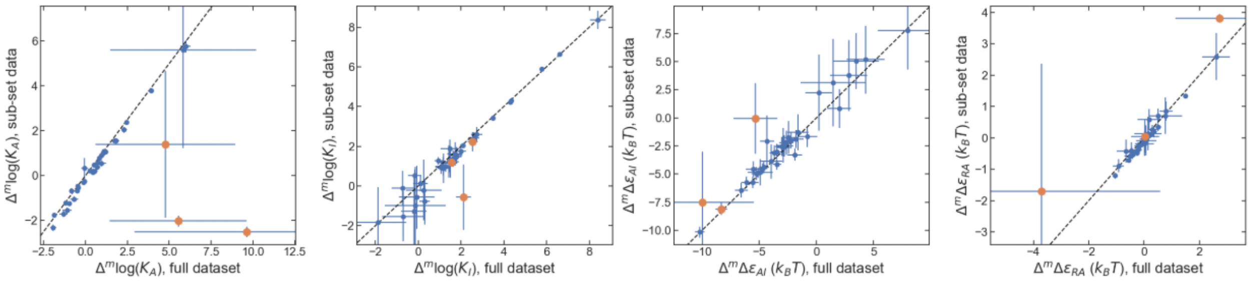


**Figure S2**. Comparison between single-operator biophysical model fit results using the full dataset (x-axis) and sub-sets of the data for each mutation (y-axis). Orange points highlight R51C, Q54R, and A82L. Plotted points are the posterior mean and error bars indicate ± one posterior standard deviation.

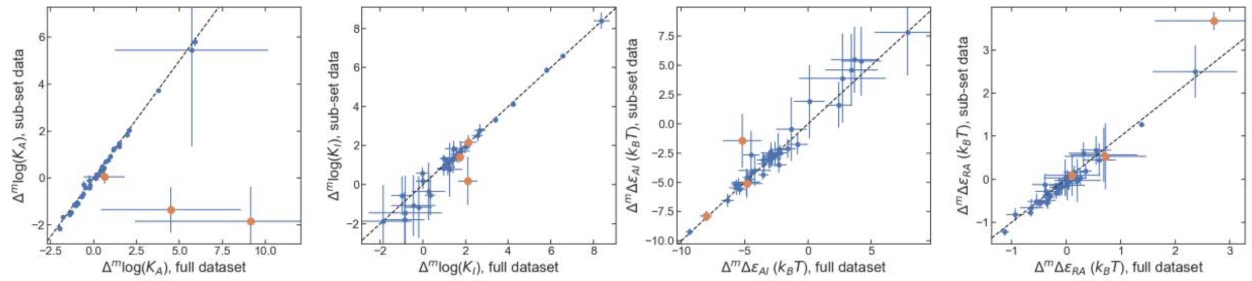


**Figure S3**. Comparison between multi-operator biophysical model fit results using the full dataset (x-axis) and sub-sets of the data for each mutation (y-axis). Orange points highlight R51C, Q54R, and A82L. Plotted points are the posterior mean and error bars indicate ± one posterior standard deviation.

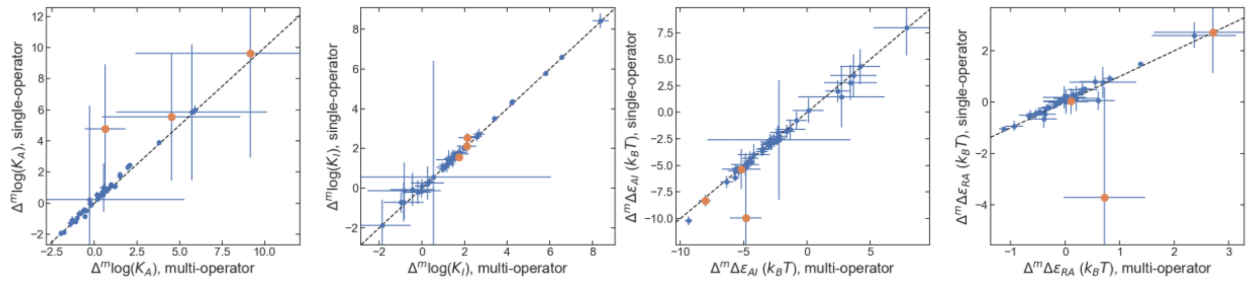


**Figure S4**. Comparison between single-operator and multi-operator biophysical models, each with the full dataset. Orange points highlight R51C, Q54R, and A82L. Plotted points are the posterior mean and error bars indicate ± one posterior standard deviation.

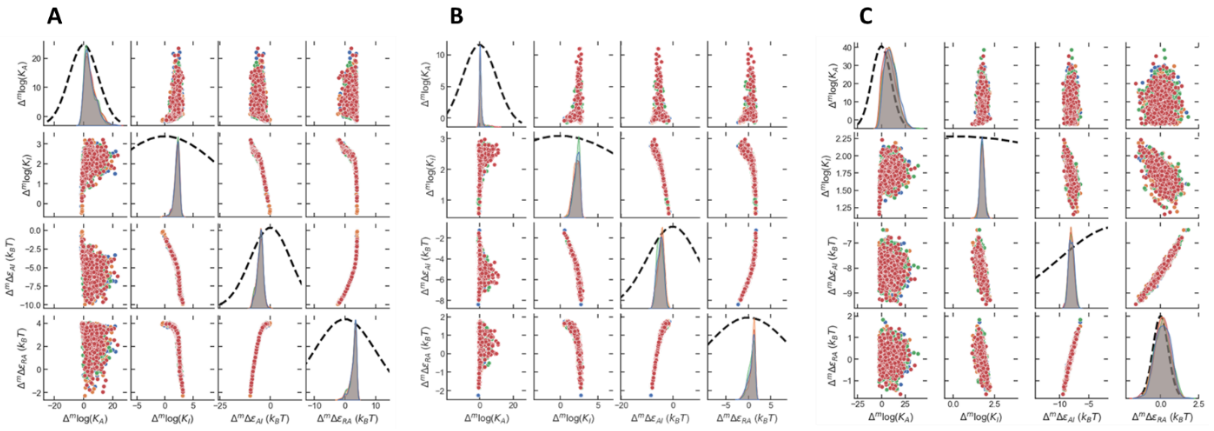


**Figure S5**. Pairplots showing samples from the posterior distribution for the mutation effects for **A:** R51C, **B:** Q54R, and **C:** A82L. Bold dashed curves in each diagonal plot are the priors distributions used with the Bayesian inference: Normal distribution with zero mean and std = 10, except for $\Delta\varepsilon_{RA}$ with mutations outside the DNA binding domain have std = 0.5. Plotted points in the scatter plots show all posterior samples, colored by chain.

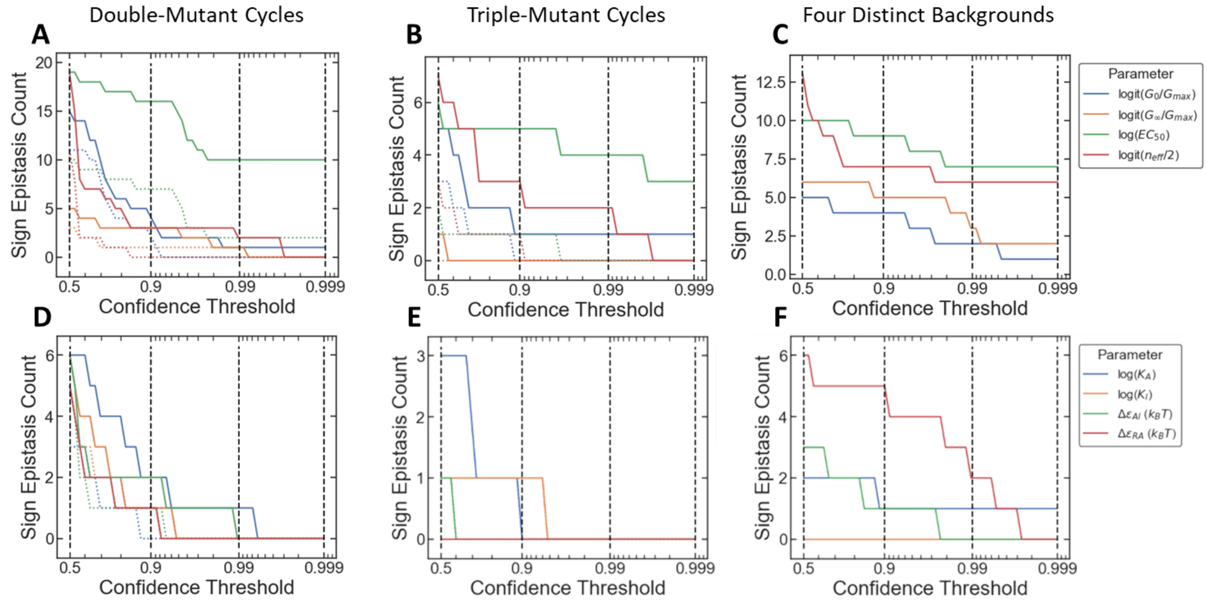


**Figure S6**. Number of mutations with sign epistasis for each phenotype parameter for **A-C:** the Hill equation model and **D-F:** the biophysical model. In **A**, **B**, **D**, and **E**, the dotted lines show results with the V136E mutation excluded.

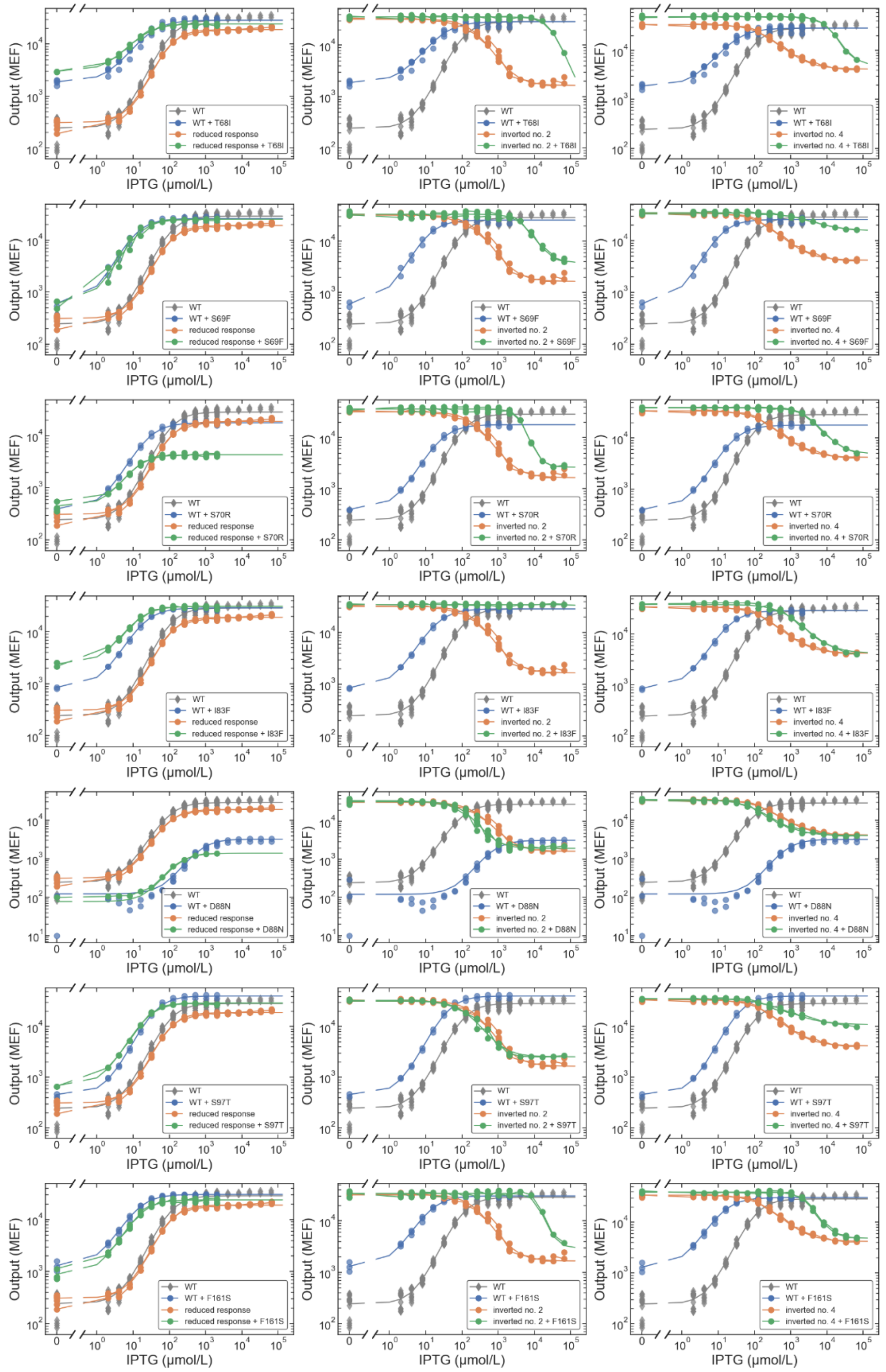


**Figure S7**. Example dose-response data for mutations with sign epistasis for the log($EC_{50}$) phenotype across the four phenotypically distinct backgrounds. Each plot shows the dose-response for the wild-type (gray), the single mutation in the wild-type background (blue), the alternate background (orange), and the mutation in the alternate background (green). Each row shows plots for a different mutation, as indicated in the legend in each plot. Individual points are shown for all measured replicates.

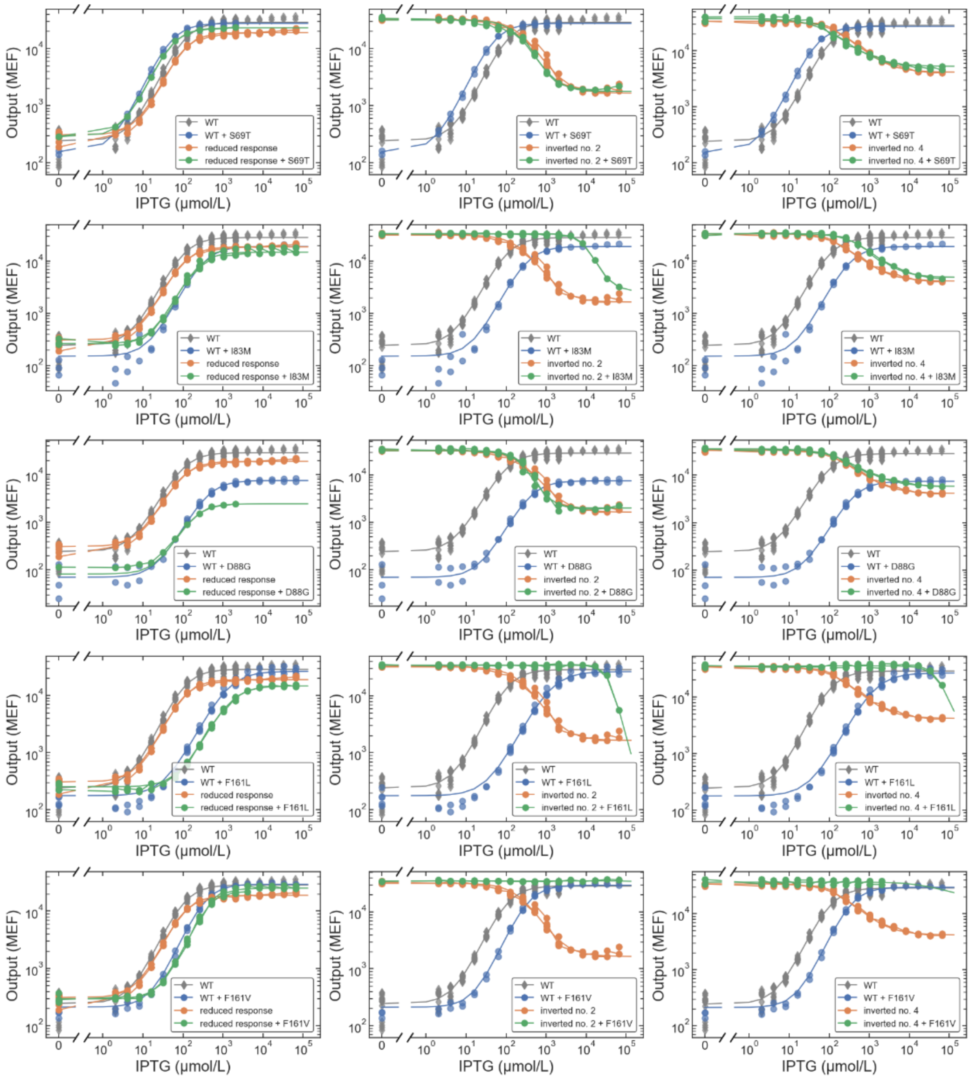


**Figure S8**. Example dose-response data for mutations at some of the same positions as those in Figure S8, but without sign epistasis for the log($EC_{50}$) phenotype. Each plot shows the dose-response for the wild-type (gray), the single mutation in the wild-type background (blue), the alternate background (orange), and the mutation in the alternate background (green). Each row shows plots for a different mutation, as indicated in the legend in each plot. Individual points are shown for all measured replicates.

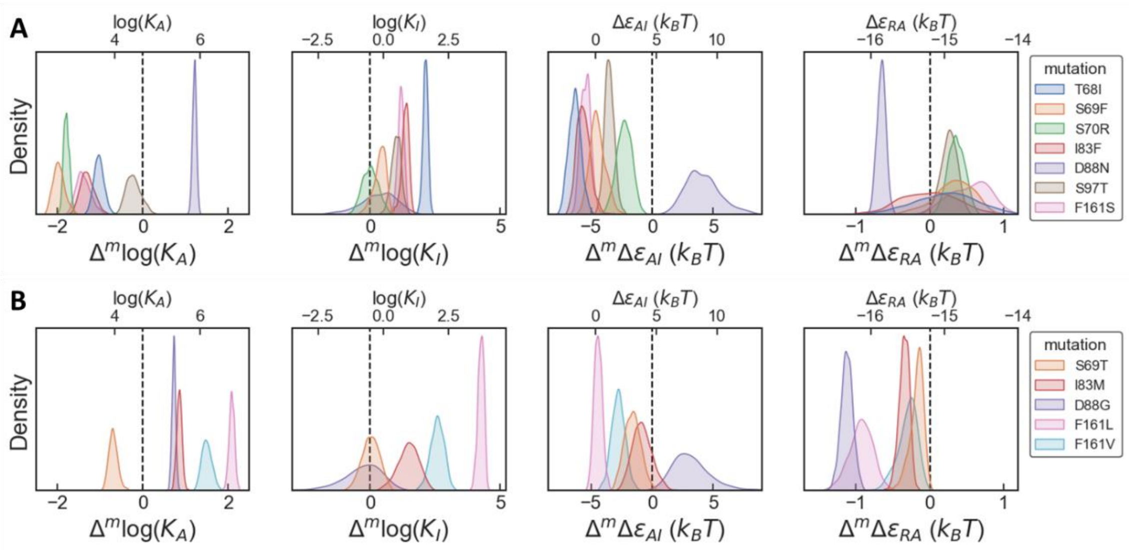


**Figure S9**. Mutation effects on the free-energy phenotypes of the biophysical model for different sets of mutations. The plots show the posterior distribution for the mutation effects from the Bayesian inference. In each plot, the bottom x-axis is the shift in the phenotype due to each mutation, and the top x-axis is the resulting phenotype value for a variant with each mutation applied to the wild-type background. The wild-type phenotypes are indicated with vertical dashed lines. **A:** Effects of mutations that have sign epistasis for the Hill model. These mutations shift the log($EC_{50}$) one direction in the wild-type and reduced-response backgrounds, but the opposite direction in the inverted backgrounds.
**B:** Effects of mutations at some of the same positions as in **A**, but that do not have sign epistasis for the Hill model.

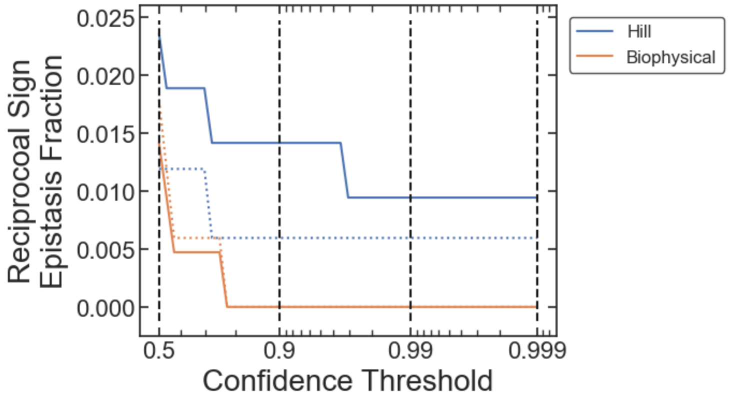


**Figure S10**. Fraction of double-mutant cycles with reciprocal sign epistasis for the Hill and biophysical models. The solid lines are the reciprocal sign epistasis fraction for all double cycles plotted vs. the confidence threshold used to determine if each mutational effect is significantly positive or negative.

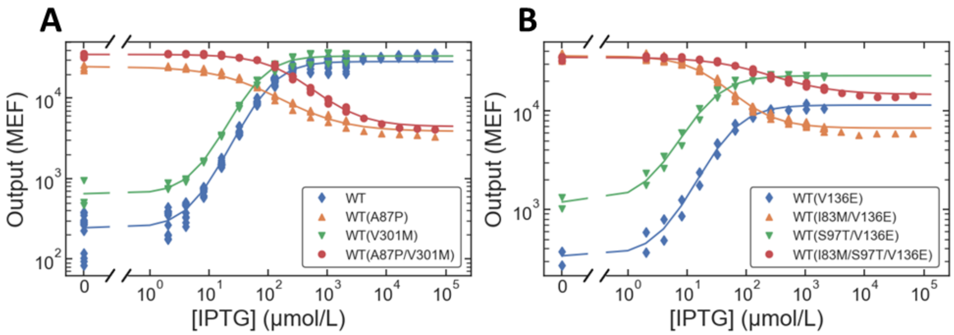


**Figure S11**. Dose-response data for double-mutant cycles with reciprocal sign epistasis for the Hill equation model. Individual points are shown for all measured replicates.

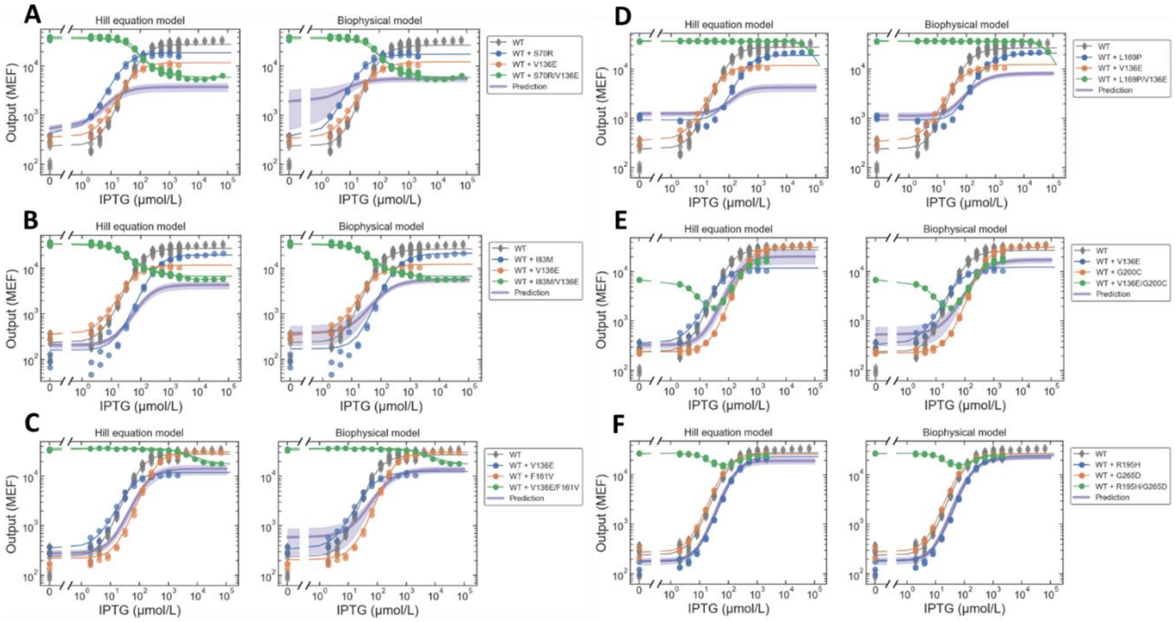


**Figure S12**. Example data and predictions for double mutants with **A-D:** inverted, or **E-F:** band-stop dose-response. Individual points are shown for all measured replicates.

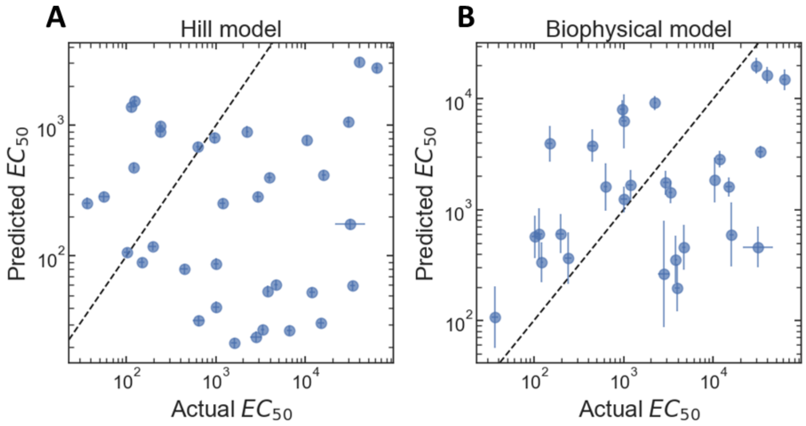


**Figure S13**. Comparison of the predictive accuracy of the Hill and biophysical models for the task of predicting the $EC_{50}$ resulting from mutations in inverted dose-response backgrounds using single-mutant effects in the wild-type background. **A:** For the Hill model, the Pearson correlation coefficient is -0.04 and the fold-RMSE is 27. **B:** For the biophysical model, the Pearson correlation coefficient is 0.44 and the fold-RMSE is 7. In each plot, plotted points are the posterior mean and error bars indicate ± one posterior standard deviation (of $\log_{10}(EC_{50})$).
