## Supplementary File S2 for "Epistasis in Allosteric Proteins: Can Biophysical Models Provide a Better Framework for Prediction and Understanding?": stan_model.hill.no_epi.stan

```
// Fit dose-response curves using Hill equation model for allosteric TFs
//     this version of the model has strictly zero epistasis
//

data {

  // Input data variables that are shared across all models 
  int<lower=1> N;        // number of data points
  vector[N] x;           // inducer concentration
  vector[N] y;           // gene expression (from cytometry) at each concentration
  vector[N] y_err;       // estimated error of gene expression at each concentration
  
  int rep[N];            // integer to indicate the measurement replicate
  int<lower=1> num_reps; // number of measurement replicates (for all variants)
  
  int<lower=1> N_contr;          // number of data points for control strains LacI deletions
  vector[N_contr] y_contr;       // gene expression (from cytometry) for control strains
  vector[N_contr] y_contr_err;   // estimated error of gene expression for control strains
  int rep_contr[N_contr];        // integer to indicate the measurement replicate for controls
  int<lower=1> num_contr_reps;   // number of measurement replicates for controls
  
  real y_max;             // geometric mean for prior on maximum gene expression value
  real g_max_prior_width; // geometric std for prior on maximum gene expression value
  
  int<lower=2> num_var;  // number of variants
  int variant[N];        // numerical index to indicate variants
  
  int<lower=1> num_mut;  // number of differrent mutations
  int<lower=0, upper=1> mut_code[num_var-1, num_mut];   // one-hot encoding for  presence of mutations in each variant; variant 0 (WT) has no mutations
  
  real rep_ratio_scale;   // parameter to set the scale for the half-normal prior on log_rep_ratio
  
  // prior for non-fluorescent control level
  real g_min_prior_mu;
  real g_min_prior_std;
  
  real rep_offset_scale;  // hyperparameter parameter to set the scale for the half-normal prior on offset_sigma

  
  // priors on wild-type Hill parameters
  real logit_g0_wt_prior_mean;
  real logit_g0_wt_prior_std;
  
  real logit_ginf_wt_prior_mean;
  real logit_ginf_wt_prior_std;
  
  real log_ec50_wt_prior_mean;
  real log_ec50_wt_prior_std;
  
  real logit_n_eff_wt_prior_mean;
  real logit_n_eff_wt_prior_std;
  
  // priors on mutational effects and epistasis for Hill parameters
  real delta_prior_width_hill;   // width of prior on delta-parameters
  
  
}

transformed data {

  // transformed data variable declarations shared by all models
  real ln_10;
  real hill_n;
  real N_NS;
  vector[19] x_out;
  real log_phi_1;
  real log_phi_2;
  real min_y;
  real g_min;           // minimum possible fluorescence (non-fluor control level)
  
  
  hill_n = 2;
  
  x_out[1] = 0;
  for (i in 2:19) {
    x_out[i] = 2^(i-2);
  }
    
  ln_10 = log(10.0);
  
  min_y = min(y);
  
  g_min = g_min_prior_mu;
  

  
}

parameters {
  // In this version of the model, the base parameters belong to the first variant (the wild-type)
  //     and there is a delta_param associated with each mutation (with additive effects)
  
  real logit_g0_wt;                 // logit-transformed g0
  vector[num_mut] logit_g0_mut;
  
  real logit_ginf_wt;               // logit-transformed ginf
  vector[num_mut] logit_ginf_mut;
  
  real log_ec50_wt;                 // log10(EC_50)
  vector[num_mut] log_ec50_mut;
  
  real logit_n_eff_wt;    // free energy for Active TF to operator
  vector[num_mut] logit_n_eff_mut;
  
  real log_g_max;                // log10 of maximum possible gene expression
  
  real<lower=0> sigma;      // scale factor for standard deviation of noise in log_y
  real<lower=0> offset_sigma;  // scale factor for standard deviation of replicate variability in g_min
  
  vector[num_reps] log_rep_ratio;              // log10 of multiplicative correction factor for different replicates
  vector[num_contr_reps] log_rep_ratio_contr;  // log10 of multiplicative correction factor for control replicates
  real<lower=0> rep_ratio_sigma;               // hyper-paramters for log_rep_ratio and log_rep_ratio_contr
  
  vector<lower=-3*rep_offset_scale, upper=3*rep_offset_scale>[num_reps] rep_offset;              // additional g_min shift for different replicates
  vector<lower=-3*rep_offset_scale, upper=3*rep_offset_scale>[num_contr_reps] rep_offset_contr;  // additional g_min shift for control replicates
  
}

transformed parameters {
  vector[num_var] g0;
  vector[num_var] ginf;
  vector[num_var] ec50;
  vector[num_var] n_eff;
  
  vector[num_var] logit_g0_var;
  vector[num_var] logit_ginf_var;
  vector[num_var] log_ec50_var;
  vector[num_var] logit_n_eff_var;
  
  vector[N] log_mean_y;
  vector[N_contr] log_mean_y_contr;
  
  // measured values with g_min subtracted
  vector[N] y_shifted;
  vector[N_contr] y_contr_shifted;
  
  logit_g0_var[1] = logit_g0_wt;
  logit_ginf_var[1] = logit_ginf_wt;
  log_ec50_var[1] = log_ec50_wt;
  logit_n_eff_var[1] = logit_n_eff_wt;
  
  g0[1] = inv_logit(logit_g0_var[1]);
  ginf[1] = inv_logit(logit_ginf_var[1]);
  ec50[1] = exp(log_ec50_var[1]);
  n_eff[1] = hill_n*inv_logit(logit_n_eff_var[1]);
  
  for (variant_num in 2:num_var) {
    logit_g0_var[variant_num] = logit_g0_wt;
    logit_ginf_var[variant_num] = logit_ginf_wt;
    log_ec50_var[variant_num] = log_ec50_wt;
    logit_n_eff_var[variant_num] = logit_n_eff_wt;
	
	for (mut in 1:num_mut) {
	  logit_g0_var[variant_num] += mut_code[variant_num-1, mut]*logit_g0_mut[mut];
	  logit_ginf_var[variant_num] += mut_code[variant_num-1, mut]*logit_ginf_mut[mut];
	  log_ec50_var[variant_num] += mut_code[variant_num-1, mut]*log_ec50_mut[mut];
	  logit_n_eff_var[variant_num] += mut_code[variant_num-1, mut]*logit_n_eff_mut[mut];
	}
	
	g0[variant_num] = inv_logit(logit_g0_var[variant_num]);
    ginf[variant_num] = inv_logit(logit_ginf_var[variant_num]);
    ec50[variant_num] = exp(log_ec50_var[variant_num]);
    n_eff[variant_num] = hill_n*inv_logit(logit_n_eff_var[variant_num]);
  }
  
  for (i in 1:N) {
	real fold_change;
	
    fold_change = g0[variant[i]] + (ginf[variant[i]] - g0[variant[i]])*(x[i]^n_eff[variant[i]])/(ec50[variant[i]]^n_eff[variant[i]] + x[i]^n_eff[variant[i]]);
    
    log_mean_y[i] = ln_10*log_g_max + log(fold_change) + log_rep_ratio[rep[i]];
	
    // measured values with g_min and rep_offset subtracted
    y_shifted[i] = y[i] - g_min - rep_offset[rep[i]];
  }
  
  for (i in 1:N_contr) {
    log_mean_y_contr[i] = ln_10*log_g_max + log_rep_ratio_contr[rep_contr[i]];
	
    y_contr_shifted[i] = y_contr[i] - g_min - rep_offset_contr[rep_contr[i]];
  }
  
}

model {
  // priors on free energy params
  logit_g0_wt ~ normal(logit_g0_wt_prior_mean, logit_g0_wt_prior_std);
  logit_ginf_wt ~ normal(logit_ginf_wt_prior_mean, logit_ginf_wt_prior_std);
  log_ec50_wt ~ normal(log_ec50_wt_prior_mean, log_ec50_wt_prior_std);
  logit_n_eff_wt ~ normal(logit_n_eff_wt_prior_mean, logit_n_eff_wt_prior_std);
  
  // priors on mutational  effects
  logit_g0_mut ~ normal(0, delta_prior_width_hill);
  logit_ginf_mut ~ normal(0, delta_prior_width_hill);
  log_ec50_mut ~ normal(0, delta_prior_width_hill);
  logit_n_eff_mut ~ normal(0, delta_prior_width_hill);
  
  // prior on max output level
  log_g_max ~ normal(log10(y_max), g_max_prior_width);
  
  // prior on scale parameter for log-normal measurement error
  sigma ~ normal(0, 1);
  
  // model of the data (dose-response curve with noise)
  y_shifted ~ lognormal(log_mean_y, sigma);
  
  // model of the control strain data (constant, max output)
  y_contr_shifted ~ lognormal(log_mean_y_contr, sigma);
  
  // prior on scale hyper-parameters for log_rep_ratio and rep_offset
  rep_ratio_sigma ~ normal(0, rep_ratio_scale);
  offset_sigma ~ normal(0, rep_offset_scale);
  
  // priors on log_rep_ratio and log_rep_ratio_contr
  log_rep_ratio ~ normal(0, rep_ratio_sigma);
  log_rep_ratio_contr ~ normal(0, rep_ratio_sigma);
  
  // priors on rep_offset, rep_offset_contr
  rep_offset ~ normal(0, offset_sigma);
  rep_offset_contr ~ normal(0, offset_sigma);
  

}

generated quantities {
  real y_out[num_var, 19];
  real fc_out[num_var, 19];
  
  real log_g0[num_var];
  real log_ginf[num_var];
  real log_ec50[num_var];
  
  real g_max;
  
  g_max = 10^log_g_max;
  
  for (variant_num in 1:num_var) {
    for (i in 1:19) {
      fc_out[variant_num, i] = g0[variant_num] + (ginf[variant_num] - g0[variant_num])*(x_out[i]^n_eff[variant_num])/(ec50[variant_num]^n_eff[variant_num] + x_out[i]^n_eff[variant_num]);
	
      y_out[variant_num, i] = g_max*fc_out[variant_num, i];
      
    }
	
	log_g0[variant_num] = log10(g0[variant_num]);
	log_ginf[variant_num] = log10(ginf[variant_num]);
	log_ec50[variant_num] = log10(ec50[variant_num]);
	
  }
  
}
```
