## Supplementary File S2 for "Epistasis in Allosteric Proteins: Can Biophysical Models Provide a Better Framework for Prediction and Understanding?": stan_model.multi-operator.no_epi.stan

```
// Fit dose-response curves to multi-operator biophysical model for allosteric TFs
//     this version of the model has strictly zero epistasis

data {

  // Input data variables that are shared across all models 
  int<lower=1> N;        // number of data points
  vector[N] x;           // inducer concentration
  vector[N] y;           // gene expression (from cytometry) at each concentration
  vector[N] y_err;       // estimated error of gene expression at each concentration
  
  int rep[N];            // integer to indicate the measurement replicate
  int<lower=1> num_reps; // number of measurement replicates (for all variants)
  
  int<lower=1> N_contr;          // number of data points for control strains LacI deletions
  vector[N_contr] y_contr;       // gene expression (from cytometry) for control strains
  vector[N_contr] y_contr_err;   // estimated error of gene expression for control strains
  int rep_contr[N_contr];        // integer to indicate the measurement replicate for controls
  int<lower=1> num_contr_reps;   // number of measurement replicates for controls
  
  real y_max;             // geometric mean for prior on maximum gene expression value
  real g_max_prior_width; // geometric std for prior on maximum gene expression value
  
  int<lower=2> num_var;  // number of variants
  int variant[N];        // numerical index to indicate variants
  
  int<lower=1> num_mut;  // number of differrent mutations
  int<lower=0, upper=1> mut_code[num_var-1, num_mut];   // one-hot encoding for  presence of mutations in each variant; variant 0 (WT) has no mutations
  
  real rep_ratio_scale;   // parameter to set the scale for the half-normal prior on log_rep_ratio
  
  // prior for non-fluorescent control level
  real g_min_prior_mu;
  real g_min_prior_std;
  
  real rep_offset_scale;  // hyperparameter parameter to set the scale for the half-normal prior on offset_sigma
  

  // Input data for priors on wild-type free energy parameters (single- and multi-operator models)
  real log_k_a_wt_prior_mean;
  real log_k_a_wt_prior_std;
  real log_k_i_wt_prior_mean;
  real log_k_i_wt_prior_std;
  real delta_eps_AI_wt_prior_mean;
  real delta_eps_AI_wt_prior_std;
  real delta_eps_RA_wt_prior_mean;
  real delta_eps_RA_wt_prior_std;
  
  // priors on mutational effects and epistasis  (single- and multi-operator models)
  real delta_prior_width;   // width of prior on "_mut" parameters (mutational effects)
  
  real<lower=0> eps_RA_prior_scale[num_mut];     // scale multiplier for width of priors on operator binding free energy term (delta_eps_RA_mut) 
  
  
  // Input data variables for multi-operator model priors
  real<lower=2> copy_num_prior_mean;  // geometric mean for prior on plasmid/operator copy number
  real<lower=0> copy_num_prior_width; // geometric std for prior on plasmid/operator copy number
  real<lower=1> R_prior_mean;         // geometric mean for prior on repressor dimer copy number
  real<lower=0> log_R_prior_width;    // geometric std for prior on repressor dimer copy number
  
}

transformed data {

  // transformed data variable declarations shared by all models
  real ln_10;
  real hill_n;
  real N_NS;
  vector[19] x_out;
  real log_phi_1;
  real log_phi_2;
  real min_y;
  real g_min;           // minimum possible fluorescence (non-fluor control level)
  
  
  hill_n = 2;
  
  x_out[1] = 0;
  for (i in 2:19) {
    x_out[i] = 2^(i-2);
  }
    
  ln_10 = log(10.0);
  
  min_y = min(y);
  
  g_min = g_min_prior_mu;
  
  N_NS = 3*4600000;
  
}

parameters {
  // In this version of the model, the base parameters belong to the first variant (the wild-type)
  //     and there is a delta_param associated with each mutation (with additive effects)

  real log_k_a_wt;         // log10 of IPTG binding affinity to active state
  vector[num_mut] log_k_a_mut;
  
  real log_k_i_wt;         // log10 of IPTG binding affinity to inactive state
  vector[num_mut] log_k_i_mut;
  
  real delta_eps_AI_wt;    // free energy difference between active and inactive states
  vector[num_mut] delta_eps_AI_mut;
  
  real delta_eps_RA_wt;    // free energy for Active TF to operator
  vector[num_mut] delta_eps_RA_mut;
  
  real log_g_max;       // log10 of maximum possible gene expression
  
  real<lower=0> sigma;      // scale factor for standard deviation of noise in log_y
  
  real<lower=0> log_copy_num;    // log10 of plasmid/operator copy number
  real<lower=0> log_R;           // log10 of repressor dimer copy number

  
  vector[num_reps] log_rep_ratio;              // log10 of multiplicative correction factor for different replicates
  vector[num_contr_reps] log_rep_ratio_contr;  // log10 of multiplicative correction factor for control replicates
  real<lower=0> rep_ratio_sigma;               // hyper-parameters for log_rep_ratio and log_rep_ratio_contr
  
  real<lower=0> offset_sigma;  // hyper-parameter for standard deviation of replicate variability in g_min
  
  vector<lower=-3*rep_offset_scale, upper=3*rep_offset_scale>[num_reps] rep_offset;              // additional g_min shift for different replicates
  vector<lower=-3*rep_offset_scale, upper=3*rep_offset_scale>[num_contr_reps] rep_offset_contr;  // additional g_min shift for control replicates
  
  
}

transformed parameters {

  vector[num_var] K_A;
  vector[num_var] K_I;
  vector[num_var] log_k_a_var;
  vector[num_var] log_k_i_var;
  vector[num_var] delta_eps_AI_var;
  vector[num_var] delta_eps_RA_var;
  
  vector[N] log_mean_y;
  vector[N_contr] log_mean_y_contr;
  
  // measured values with g_min subtracted
  vector[N] y_shifted;
  vector[N_contr] y_contr_shifted;

  real N_S;
  real R;

  N_S = 10^log_copy_num;
  R = 10^log_R;
  
  
  log_k_a_var[1] = log_k_a_wt;
  log_k_i_var[1] = log_k_i_wt;
  delta_eps_AI_var[1] = delta_eps_AI_wt;
  delta_eps_RA_var[1] = delta_eps_RA_wt;
  
  K_A[1] = 10^log_k_a_var[1];
  K_I[1] = 10^log_k_i_var[1];
  
  for (variant_num in 2:num_var) {
    log_k_a_var[variant_num] = log_k_a_wt;
    log_k_i_var[variant_num] = log_k_i_wt;
    delta_eps_AI_var[variant_num] = delta_eps_AI_wt;
    delta_eps_RA_var[variant_num] = delta_eps_RA_wt;
	
	for (mut in 1:num_mut) {
	  log_k_a_var[variant_num] += mut_code[variant_num-1, mut]*log_k_a_mut[mut];
	  log_k_i_var[variant_num] += mut_code[variant_num-1, mut]*log_k_i_mut[mut];
	  delta_eps_AI_var[variant_num] += mut_code[variant_num-1, mut]*delta_eps_AI_mut[mut];
	  delta_eps_RA_var[variant_num] += mut_code[variant_num-1, mut]*delta_eps_RA_mut[mut];
	}
	
	K_A[variant_num] = 10^log_k_a_var[variant_num];
    K_I[variant_num] = 10^log_k_i_var[variant_num];
  }
  
  
  for (i in 1:N) {
    real c1;
    real c2;
    real pA;
	real xRA;
	real lam;
	real fold_change;
	
    c1 = (1 + x[i]/K_A[variant[i]])^hill_n;
    c2 = ( (1 + x[i]/K_I[variant[i]])^hill_n ) * exp(-delta_eps_AI_var[variant[i]]);
	pA = c1/(c1+c2);
	xRA = exp(-delta_eps_RA_var[variant[i]]);
	
	lam = -N_NS + pA*R - N_S*xRA + pA*R*xRA;
    lam = lam + sqrt(4*pA*R*xRA*(N_NS + N_S - pA*R) + (N_NS + N_S*xRA - pA*R*(1 + xRA))^2);
    lam = lam/(2*xRA*(N_NS + N_S - pA*R));
	
    fold_change = 1/(1 + lam*xRA);
	
    log_mean_y[i] = ln_10*log_g_max + log(fold_change) + log_rep_ratio[rep[i]];
	
    // measured values with g_min and rep_offset subtracted
    y_shifted[i] = y[i] - g_min - rep_offset[rep[i]];
  }
  
  for (i in 1:N_contr) {
    log_mean_y_contr[i] = ln_10*log_g_max + log_rep_ratio_contr[rep_contr[i]];
	
    y_contr_shifted[i] = y_contr[i] - g_min - rep_offset_contr[rep_contr[i]];
  }
  
  
}

model {

  // priors on free energy params
  log_k_a_wt ~ normal(log_k_a_wt_prior_mean, log_k_a_wt_prior_std);
  log_k_i_wt ~ normal(log_k_i_wt_prior_mean, log_k_i_wt_prior_std);
  delta_eps_AI_wt ~ normal(delta_eps_AI_wt_prior_mean, delta_eps_AI_wt_prior_std);
  delta_eps_RA_wt ~ normal(delta_eps_RA_wt_prior_mean, delta_eps_RA_wt_prior_std);
  
  log_k_a_mut ~ normal(0, delta_prior_width/ln_10); // factor of 1/ln_10 is to compensate for use of log10 instead of ln
  
  log_k_i_mut ~ normal(0, delta_prior_width/ln_10); // factor of 1/ln_10 is to compensate for use of log10 instead of ln
  
  delta_eps_AI_mut ~ normal(0, delta_prior_width);
  
  for (mut in 1:num_mut) {
    delta_eps_RA_mut[mut] ~ normal(0, delta_prior_width*eps_RA_prior_scale[mut]);
  }
  
  // prior on max output level
  log_g_max ~ normal(log10(y_max), g_max_prior_width);
  
  // prior on scale parameter for log-normal measurement error
  sigma ~ normal(0, 1);
  
  // model of the data (dose-response curve with noise)
  y_shifted ~ lognormal(log_mean_y, sigma);
  
  // model of the control strain data (constant, max output)
  y_contr_shifted ~ lognormal(log_mean_y_contr, sigma);

  
  // prior on scale hyper-parameters for log_rep_ratio and rep_offset
  rep_ratio_sigma ~ normal(0, rep_ratio_scale);
  offset_sigma ~ normal(0, rep_offset_scale);
  
  // priors on log_rep_ratio and log_rep_ratio_contr
  log_rep_ratio ~ normal(0, rep_ratio_sigma);
  log_rep_ratio_contr ~ normal(0, rep_ratio_sigma);
  
  // priors on rep_offset, rep_offset_contr
  rep_offset ~ normal(0, offset_sigma);
  rep_offset_contr ~ normal(0, offset_sigma);
  
  
  // priors on plasmid/operator and repressor dimer copy numbers
  log_copy_num ~ normal(log10(copy_num_prior_mean), copy_num_prior_width);
  log_R ~ normal(log10(R_prior_mean), log_R_prior_width);


}

generated quantities {

  real g_max;
  real y_out[num_var, 19];
  real fc_out[num_var, 19];
  
  g_max = 10^log_g_max;
  
  
  for (variant_num in 1:num_var) {
    for (i in 1:19) {
      real c1;
      real c2;
      real pA;
	  real xRA;
	  real lam;
	  real f_c;
	  
      c1 = (1 + x_out[i]/K_A[variant_num])^hill_n;
      c2 = ( (1 + x_out[i]/K_I[variant_num])^hill_n ) * exp(-delta_eps_AI_var[variant_num]);
	  pA = c1/(c1+c2);
	  xRA = exp(-delta_eps_RA_var[variant_num]);
	
	  lam = -N_NS + pA*R - N_S*xRA + pA*R*xRA;
      lam = lam + sqrt(4*pA*R*xRA*(N_NS + N_S - pA*R) + (N_NS + N_S*xRA - pA*R*(1 + xRA))^2);
      lam = lam/(2*xRA*(N_NS + N_S - pA*R));
	
      f_c = 1/(1 + lam*xRA);
	
      y_out[variant_num, i] = g_max*f_c;
      fc_out[variant_num, i] = f_c;
    }
  }
  

  
}
```
